# BK channels have opposite effects on sodium versus calcium-mediated action potentials in endocrine pituitary cells

**DOI:** 10.1101/477976

**Authors:** Geir Halnes, Simen Tennøe, Trude M. Haug, Gaute T. Einevoll, Finn-Arne Weltzien, Kjetil Hodne

## Abstract

Pituitary endocrine cells fire action potentials (APs) to regulate their cytosolic Ca^2+^ concentration and hormone secretion rate. Depending on animal species, cell type, and biological conditions, pituitary APs are generated either by TTX-sensitive Na^+^ currents (*I*_*Na*_), high-voltage activated Ca^2+^ currents (*I*_*Ca*_), or by a combination of the two. Previous computational models of pituitary cells have mainly been based on data from rats, where *I*_*Na*_ is largely inactivated at the resting potential, and spontaneous APs are exclusively mediated by *I*_*Ca*_. As a part of the previous modeling studies, a paradoxical role was identified for the big conductance K^+^ current (*I*_*BK*_), which was found to prolong the duration of *I*_*Ca*_-mediated APs, and sometimes give rise to pseudo-plateau bursts, contrary to what one would expect from a hyperpolarizing current. Unlike in rats, spontaneous *I*_*Na*_-mediated APs are consistently seen in pituitary cells of several other animal species, including several species of fish. In the current work we develop the, to our knowledge, first *computational model* of a pituitary cell that fires *I*_*Na*_-mediated APs. Although we constrain the model to experimental data from gonadotrope cells in the teleost fish medaka (*Oryzias latipes*), it may likely provide insights also into other pituitary cell types that fire *I*_*Na*_-mediated APs. In the current work, we use the model to explore how the effect of *I*_*BK*_ depends on the AP generating mechanisms of pituitary cells. We do this by comparing simulations on the medaka gonadotrope model (two versions thereof) with simulations on a previously developed model of a rat pituitary cell. Interestingly, we find that *I*_*BK*_ has the opposite effect on APs in the two models, i.e. it reduces the duration of already fast *I*_*Na*_-mediated APs in the medaka model, and prolongs the duration of already slow *I*_*Ca*_-mediated APs in the rat model.

**Author summary:** Excitable cells elicit electrical pulses called action potentials (APs), which are generated and shaped by a combination of ion channels in the cell membrane. While neurons use APs for interneuronal communication and heart cells use them to generate heart-beats, pituitary cells use APs to regulate their cytosolic Ca^2+^ concentration, which in turn controls their hormone secretion rate. The amount of Ca^2+^ that enters the pituitary cell during an AP depends strongly on how long it lasts, and it is therefore important to understand the mechanisms that control this. Depending on animal species and biological conditions, pituitary APs may be initiated either by Ca^2+^ channels or Na^+^ channels. Here, we explore the differences between the two scenarios by comparing simulations on two different computer models: (i) a previously developed model which fires Na^+^-based APs, adapted to data from pituitary cells in rats, and (ii) a novel model that fires Ca^2+^-based APs, adapted to data from pituitary cells in the fish medaka. Interestingly, we find that the role of big conductance K^+^ (BK) channels, which are known to affect the duration of the AP, are opposite in the two models, i.e., they act to prolong Ca^2+^-based APs while they act to shorten Na^+^-based APs.

## 1 Introduction

The electrodynamics of excitable cells is generated by a combination of ion channels in the plasma membrane, which are typically characterized by their voltage and/or Ca^2+^ dependence. While neurons primarily use action potentials as a means of interneuronal communication, and cardiac cells use them to generate heartbeats, the primary role of APs in endocrine pituitary cells is to regulate the cytosolic Ca^2+^ concentration, which in turn controls the hormone secretion rate in these cells [1]. Hormone secretion often occurs as a response to hormonal stimuli from the hypothalamus, peripheral endocrine glands, and other types of pituitary cells. However, many endocrine cells are also spontaneously active [1–10]. The spontaneous activity is partly a means to regulate the re-filling of intracellular Ca^2+^ stores, but in several cells also leads to a basal release of hormones. An understanding of the mechanisms regulating the electrodynamics of these cells is therefore fundamental for understanding their overall functioning.

While neuronal APs are are predominantly mediated by TTX-sensitive Na^+^ currents (*I*_*Na*_), AP generation in endocrine cells depends strongly on high-voltage-activated Ca^2+^ currents (*I*_*Ca*_), which in addition to their role in affecting the voltage dynamics of the cell, also are the main source of Ca^2+^ entry through the plasma membrane [3, 11, 12]. In some studies of endocrine cells, APs were exclusively mediated by *I*_*Ca*_, and the spontaneous membrane excitability was insensitive or nearly so to TTX [1, 2, 13–16]. In other studies, APs were evoked by a combination of *I*_*Ca*_ and *I*_*Na*_ [4, 7, 17, 18]. The strong involvement of *I*_*Ca*_ could explain why pituitary APs typically last longer (typically some tens of milliseconds [8]) than neuronal APs (a few milliseconds), which are mainly mediated by *I*_*Na*_.

All endocrine cells express *I*_*Na*_ [8], and TTX sensitive APs can typically be triggered by current injections from hyperpolarized holding potentials even in cells where they are not elicited spontaneously [4, 17, 19, 20]. The reason why the spontaneous activity is TTX insensitive is likely that a major fraction of *I*_*Na*_ is inactivated at the resting membrane potential [15, 16]. The reason why this is not always the case, may be that the resting potentials vary greatly between different studies. Only for rat somatotropes, resting potentials ranging as wide as from −30 mV [13] to −80 mV [18] have been reported. In cells with hyperpolarized resting potentials (−80 mV), TTX was found to block single, brief action potentials, while action potentials of long duration and low amplitude persisted [18], indicating that both a *I*_*Ca*_ and *I*_*Na*_ component were present and that the two had different time-courses. However, the more typical resting potentials for rat pituitary cells lie in the range between *−*50 mV and *−*60 mV, and at these resting levels, *I*_*Na*_ tends to be inactivated and the spontaneous activity TTX insensitive (see reviews in [8, 21]).

Computational models constructed to capture the essential activity of pituitary cells have predominantly relied on rat (or mouse) data, and have not included *I*_*Na*_ [3, 9, 22–27]. When *I*_*Na*_ was included in a recent modeling study, its role was mainly in modulating the firing patterns but was not essential for AP firing as such [28]. The main focus of these models was thus on exploring the interplay between the depolarizing *I*_*Ca*_ and various K^+^ currents responsible for shaping the repolarization following an AP. The essentials of this interplay were nicely captured in a relatively simple computational model by Tabak et al. [9]. In this model, the AP upstroke was exclusively mediated by *I*_*Ca*_, while the repolarizing phase was mediated by three K^+^ currents (*I*_*K*_, *I*_*BK*_ and *I*_*SK*_). The model elicited spontaneous APs with a duration of about 40 ms, which is quite representative for what is seen experimentally in rat pituitary cells. In addition, the model was able to explain the important and paradoxical role that the big conductance Ca^2+^-activated K^+^ current (*I*_*BK*_) has in rat endocrine cells, where it was found to make APs broader. The explanation to this broadening effect, which is the opposite of what one would expect from a hyperpolarizing current [23], was that *I*_*BK*_ reduces the AP amplitude, and thereby prevents the (high-threshold) activation of the delayed rectifying K^+^ current (*I*_*K*_), which otherwise would become a more powerful hyperpolarizing current [9, 23]. By inhibiting *I*_*K*_, *I*_*BK*_ indirectly becomes facilitating, and leads to broader APs and sometimes to voltage plateaus and so-called *pseudo plateau bursts* [8, 23], which are believed to be more efficient than regular APs in evoking hormone secretion (see e.g. [29]). By manipulating the BK expression experimentally and in computational models, it was further shown that the difference between bursty endocrine cell types such as somatotropes and lactotropes, and regularly spiking cell types such as gonadotropes and corticotrophs, could be explained by the different cell types having different levels of *I*_*BK*_ expression [9, 28].

The modeling studies cited above were predominantly based on data from rats, and there are reasons to believe that teleost pituitary cells are different in terms of their dynamical properties. Firstly, TTX-sensitive spontaneous activity has been seen in goldfish resting at *−*60 mV [4], and TTX sensitive APs has been evoked from a holding potential as high as −50 mV in pituitary cells in cod [7], suggesting that *I*_*Na*_ may be more available in resting pituitary cells in fish [4]. Secondly, data from goldfish [4] and tilapia [5] indicate that the AP duration is shorter in fish (*<* 10 ms) compared to rats (several tens of ms), and also this could indicate a stronger involvement of *I*_*Na*_. A third difference between fish and rat pituitary cells is in the role of *I*_*BK*_. *I*_*BK*_ is almost absent in rat gonadotropes [23], and this was proposed as an explanation to why these cells tend to be less bursty than other pituitary cell types [1, 9, 28]. In contrast, *I*_*BK*_ is highly expressed in medaka gonadotropes, but without making these cells bursty [12]. The indication that there are differences between rat and fish pituitary cells are supported by experiments presented in the current work, performed on gonadotrope cells in medaka. We show that these cells elicit brief spontaneous APs that (unlike spontaneous APs in rats) to a large degree are mediated by TTX sensitive Na^+^ currents (*I*_*Na*_). Furthermore, we show that *I*_*BK*_ acts to make APs narrower in medaka gonadotropes, and thus have the opposite effect of what they do in rat pituitary cells.

Since previous computational models based on rat data seem unsuited to describe the spontaneous activity of fish pituitary cells, we here present novel pituitary cell models constrained to data from luteinizing hormone-producing medaka gonadotropes. The main, and more general, aim of this study is to use computational modeling to explore the differences between pituitary cells that fire APs exclusively mediated by *I*_*Ca*_ (like the previous rat models) and pituitary cells that fire APs that are predominantly mediated by *I*_*Na*_ (like the here developed medaka models), with a special focus on the role that *I*_*BK*_ has in the two cases. To do so, we compare simulations run on three different computational models:

1. **RAT**. The first model, which we refer to as RAT, is a reproduced version of the model by Tabak et al. [9].
2. **MEDAKA 1**. In the second model, which we refer to as MEDAKA 1, *I*_*Ca*_ from RAT is replaced with a pair of depolarizing currents *I*_*Na*_ and *I*_*Ca*_ with kinetics constrained to voltage-clamp data from gonadotrope cells in medaka. In MEDAKA 1, all other currents were kept identical to that in RAT, as this allowed us to make a direct comparison between two models where only the AP generating mechanisms were different.
3. **MEDAKA 2**. In the third model, which we refer to as MEDAKA 2, *I*_*Na*_ and *I*_*Ca*_ were kept as in MEDAKA 1, but the remaining set of ionic currents were adjusted in order to obtain a model that better replicates the essential response features of gonadotrope cells in medaka.

By simulations, we show that MEDAKA 1 and MEDAKA 2 produce spontaneous APs that are faster than those in RAT, and thus more suited to describe the firing properties of fish pituitary cells. We further show that *I*_*BK*_ has the opposite effect in the medaka models from what it has in RAT, and suggest that *I*_*BK*_ acts as a mechanisms that makes slow (*I*_*Ca*_-generated) pituitary APs broader, and fast (*I*_*Na*_-generated) pituitary APs briefer. To our knowledge, MEDAKA 1 and MEDAKA 2 are the first computational models that describe *I*_*Na*_-based APs in endocrine cells. Although the models were tailored to represent gonadotrope cells in medaka, we believe that they are of a more general value for improving our understanding of *I*_*Na*_-based APs in the pituitary, which are elicited by several endocrine cell types and in several animal species, depending on biological conditions [4, 7, 17, 18, 30–32].

## Results

### Characteristic response patterns of medaka gonadotropes

The general electrophysiological properties of gonadotrope cells in medaka were assessed through a series of voltage clamp and current clamp experiments. Voltage clamp experiments used to develop kinetics models of Na^+^ and Ca^2+^ currents in MEDAKA 1 and MEDAKA 2 are presented in the Methods section. Here, we focus on the key properties of spontaneous APs as recorded by current clamp. Selected, representative experiments are shown in Fig 1.

**Fig 1.**
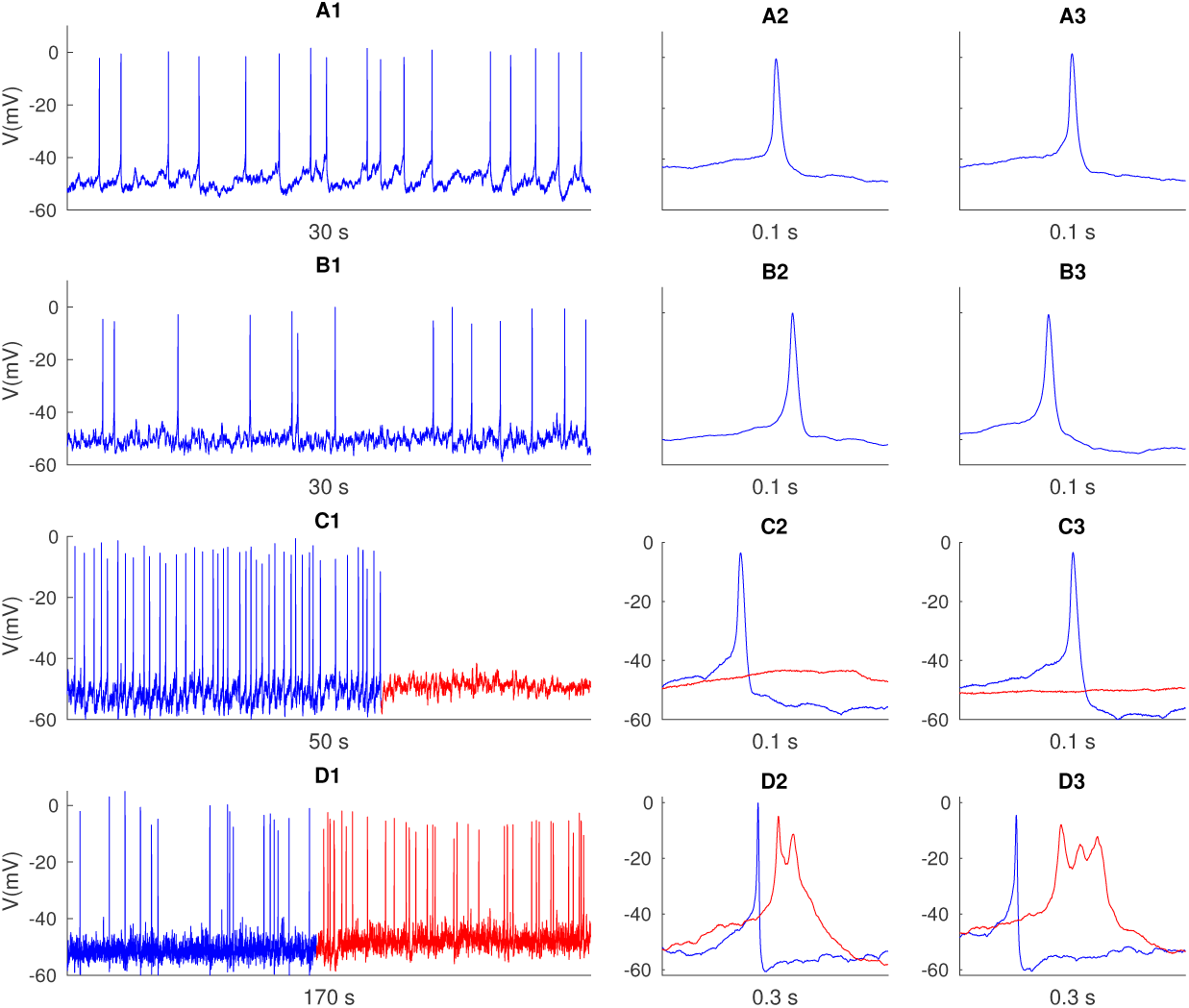
Experimental voltage recordings. (A1-B1) Spontaneous AP firing in two selected cells. (A2-A3) Close-ups of selected APs from the cell in A1. (B2-B3) Close-ups of selected APs from the cell in A2. (C1) Spontaneous activity before (blue) and after (red) TTX application. (C2-C3) Close-ups of two selected events before (blue) and after (red) TTX application. (D1) Spontaneous activity before (blue) and after (red) paxilline application. (D2-D3) Close-ups of two selected events before (blue) and after (red) paxilline application. The firing rates in the various recordings were 0.64 Hz (A1), 0.57 Hz (B1), 1.22 Hz (C1, before TTX), 0.17 Hz (D1, before paxilline) and about 0.35 Hz (D1, after paxilline). AP durations varied between 3 and 7 ms, with mean durations of 3.7 ms (A1), 4.9 ms (B1), 3.7 ms (C1, before TTX). In (D1), average AP durations were 4.2 ms before paxilline, and 25 ms after paxilline. Mean AP peak values were −0.4 mV (A1), −3.4 mV (B1), −5.1 mV (C1, before TTX), −3.1 mV (D1, before paxilline), and −6.4 mV (D1, after paxilline). AP width was calculated at half max amplitude between −50 mV and AP peak. The experiments were performed on gonadotrope luteinizing hormone-producing cells in medaka. All depicted traces were corrected with a liquid junction potential of −9 mV. The time indicated below each panel refers to the duration of the entire trace shown.

Although variations were observed, the medaka gonadotropes typically had a resting potential around −50 mV, which is within the range found previously for goldfish [4]and cod [10] gonadotropes. As for goldfish gonadotropes, the majority of medaka gonadotropes fired spontaneous APs, and with peak voltages slightly below 0 mV. The spontaneous APs were always regular spikes (i.e., not bursts) and typically had an average duration between 3 and 7 ms (blue traces in Fig 1). Similar brief AP waveforms have been seen in previous studies on fish [4, 5, 19], while the APs reported for rat gonadotrope cells are typically slower, i.e. from 10-100 ms [8]. The spontaneous AP activity was completely abolished by TTX application (Fig 1B), as has previously also been seen for goldfish somatotropes [19].

Finally, we explored how paxilline (an *I*_*BK*_ blocker) affected the spontaneous activity of gonadotrope medaka cells. In the experiment shown in Fig 1D, paxilline increased the firing rate and slightly reduced the mean AP peak amplitude, but these effects were not seen consistently in experiments using paxilline application. However, in all experiments, paxilline application was followed by a small increase of the resting membrane potential (Fig 1C), and a broadening of the AP waveform (Fig 1D2-D3). Similar effects have been seen in goldfish somatotropes, where application of tetraethylammonium (a general blocker of Ca^2+^ gated K^+^ currents) lead to broadening of APs [19]. The effect of *I*_*BK*_ in goldfish and medaka gonadotropes is thus to make APs narrower, which is the opposite of what was found in rat pituitary cells, where *I*_*BK*_ lead to broader APs and sometimes to burst-like activity [9, 23].

### Computational models of pituitary cells

In this work, we have compared simulations performed on three different pituitary cell models. The first model, RAT, is a replication of the previously published model by Tabak et al. [9]. This model was originally adapted to electrophysiological data from rat pituitary cells. As summarized by Eq 1, RAT contains a leakage current *I_leak_*, three K^+^ currents *I*_*K*_, *I*_*BK*_ and *I*_*SK*_, and a Ca^2+^ current 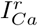. In addition, a gaussian noise stimulus was added in selected simulations (see Methods).

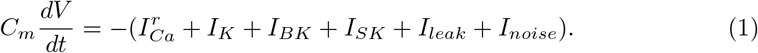

The two other models are novel for this work and are the to date first pituitary cell models based on teleost data, and the first to elicit APs that are predominantly mediated by Na^+^ currents. In these models, the Ca^2+^ current 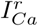 from RAT was replaced by a pair of depolarizing currents, i.e., a novel Ca^2+^ current 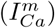 and a Na^+^ current (*I*_*Na*_), which were adapted to voltage clamp data from gonadotrope cells in medaka (see Methods). We present two versions of the medaka model, both of which are summarized by Eq 2:

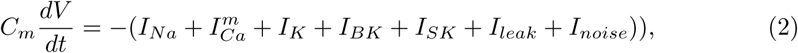

In the first version, MEDAKA 1, the three K^+^ currents *I*_*K*_, *I*_*BK*_ and *I*_*SK*_ were identical to those in RAT, so that only the depolarizing, AP generating mechanisms were different. In the second teleost model, MEDAKA 2, we adjusted *I*_*K*_, *I*_*BK*_ and *I*_*SK*_ so that the model had an AP shape and AP firing rate that were in better agreement with the experimental data in Fig. 1. By comparing RAT to MEDAKA 1, we could then explore the difference between a model with Ca^2+^ APs and one with Na^+^/Ca^2+^ APs with all other mechanisms being the same. By comparing RAT to MEDAKA 2, we could explore the difference two models that were more representative for experimental data from rat versus medaka. The differences between RAT, MEDAKA 1 and MEDAKA 2 are summarized in Table 1, which lists all the parameter values that were not identical in all three models, and Fig. 2 which shows the kinetics of all included ion channels.

**Table 1.** Model parameters differing between the models RAT, MEDAKA 1 and MEDAKA 2. *g*_*BK*_ was varied between simulations, and had values between 0 and the (maximum) value given the table in the respective models. * 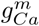 had the units of a permeability (see Methods subsection titled “Models” for explanation). *\*\**A justification for using different values for *E*_*leak*_ is given in the Methods-section.

|  | RAT | MEDAKA 1 | MEDAKA 2 | Units |
| --- | --- | --- | --- | --- |
| $g_{Ca}^r$ | 0.64 | 0 | 0 | mS/cm<sup>2</sup> |
| $g_{Ca}^m$ | 0 | 0.2 | 0.2 | cm/s* |
| $g_{Na}$ | 0 | 70 | 70 | mS/cm<sup>2</sup> |
| $g_K$ | 0.96 | 0.96 | 1.3 | mS/cm<sup>2</sup> |
| $g_{BK}$ | 0.32 | 0.32 | 1.26 | mS/cm<sup>2</sup> |
| $g_{SK}$ | 0.32 | 0.32 | 0.96 | mS/cm<sup>2</sup> |
| $\tau_K$ | 30 | 30 | 5 | ms |
| $v_f$ | -20 | -20 | -15 | mV |
| $E_{leak}^{**}$ | -50 | -45 | -45 | mV |

**Fig 2.**
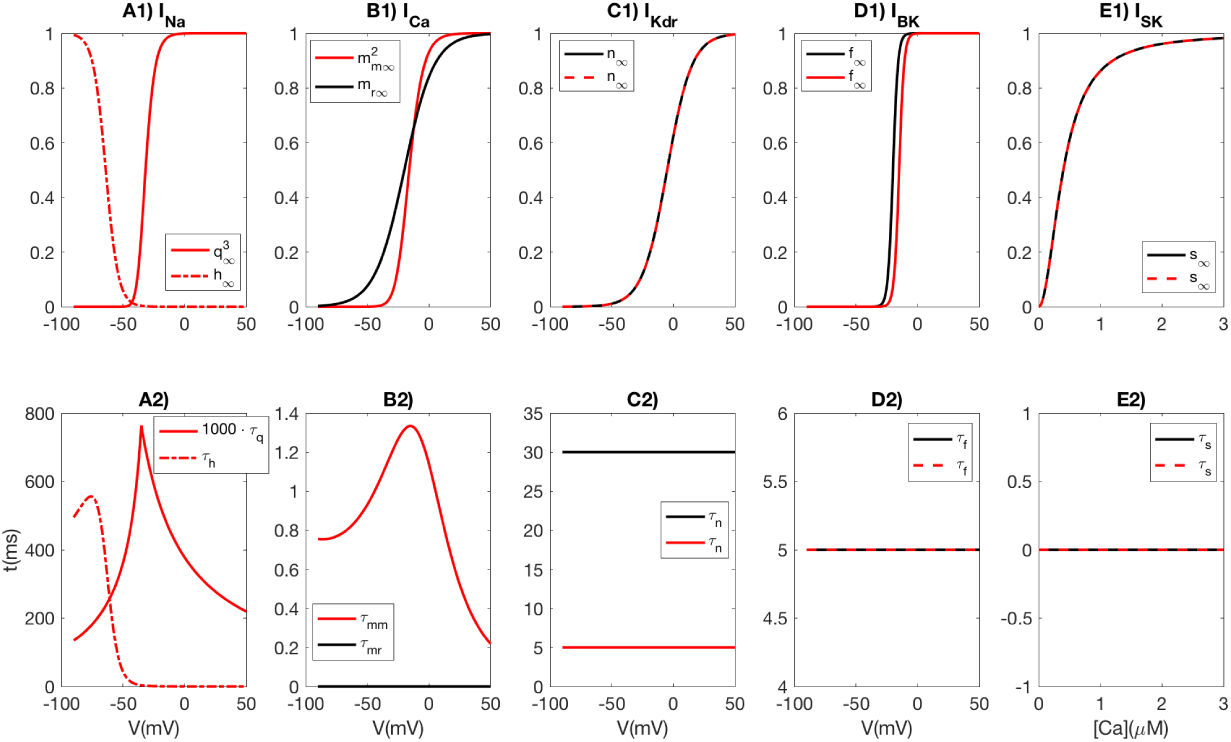
Ion channel kinetics. (A1) *I*_*Na*_ had three activation variables (*q*) and one inactivation variable (*h*). (B1) 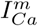 had two activation variables (*m*) in MEDAKA 1 and 2, and 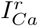 had one activation variable (*m*) in RAT. (C1) *I*_*K*_ had one activation variable (*n*). (D1) *I*_*BK*_, had one activation variable (*f*). (E1) *I*_*SK*_ was Ca^2+^ activated with one activation variable *s*. (A2-B2) Voltage dependent activation time constants were determined for *I*_*Na*_ (A2) and 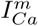 (red curve in B2) used in MEDAKA 1 and MEDAKA 2. Voltage-independent activation-time constants were used for *I*_*K*_ (C2), *I*_*BK*_ (D2) and *I*_*SK*_ (E2). (A-E) Black curves denote the RAT model, while red curves denote MEDAKA 2. MEDAKA 1 had the same kinetics as MEDAKA 2 for *I*_*Na*_ and 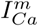 (A-B), and the same as RAT for the three K^+^ currents (C-E).

For a full description of the models, we refer to the Methods section, but a brief overview is given here. *I*_*Na*_ activated in the range between *−*50 mV and *−*10 mV, with half activation at *−*32 mV (Fig. 2A1), quite similar to what was previously found in goldfish gonadotropes [4]. *I*_*Na*_ inactivated in the range between *−*90 mV and *−*40 mV, with half-inactivation at *−*64 mV, which was lower than in goldfish, where the half-inactivation was found to be around *−*50 mV [4]. With the activation kinetics adapted to medaka data, only 6 % of the *I*_*Na*_ was available at the typical resting potential of *−*50 mV. The fact that medaka still showed TTX-sensitive spontaneous activity thus suggests that *I*_*Na*_ is very highly expressed in these cells.

Both *I*_*Na*_ and 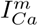 had fast activation, *I*_*Na*_ being slightly faster with a time constant of about 0.5 - 0.8 ms in the critical voltage range (Fig. 2A2), whereas *I*_*Ca*_ had a time constant *>*1 ms in the critical voltage range (Fig. 2B2). 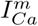 activated in the range between *−*40 mV and +10 mV, with a half activation at 16 mV (red curve in Fig. 2B1). This activation curve was much steeper than 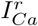 in RAT (black curve in Fig. 2B1), which was based on data from rat lactotropes [33]. The high activation threshold suggests that 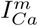 is unsuitable for initiating spontaneous APs in medaka gonadotropes, making spontaneous activity critically dependent on *I*_*Na*_.

The remaining currents (*I*_*K*_, *I*_*BK*_ and *I*_*SK*_) were all adopted from the RAT model by Tabak et al. [9], using simplified kinetics descriptions with voltage independent time constants. As explained above, these were used in their original form in RAT and MEDAKA 1, but were adapted in MEDAKA 2.

### BK currents have opposite effects on Na^*+*^-versus Ca^*2+*^-mediated action potentials

It has previously been shown that *I*_*BK*_ acts to broaden APs and promote bursting in rat pituitary cells [9, 23, 28]. In contrast, a high *I*_*BK*_ expression in medaka gonadotropes [12] does not make these cells bursty. On the contrary, the experiments in Fig. 1D showed that medaka gonadotropes became more bursty when BK channels were blocked. We hypothesized that opposite effects of BK channels in rats versus medaka are not due to the BK channels being different, but rather due to (the same) BK channels being activated in different ways in the two systems.

Here, we have used computational modeling to test this hypothesis, and have compared simulations on the models RAT, MEDAKA 1 and MEDAKA 2 for different levels of *I*_*BK*_ expression (Fig. 3). Following the definition used by Tabak et al. [9], plateau-proceeded broad APs of duration longer than 60 ms were defined as bursts (see Methods). When BK was fully expressed in the models (i.e. had the values for *g*_*BK*_ given in Table 1), MEDAKA 1 and MEDAKA 2 fired relatively narrow APs that were not classified as bursts, while RAT consistently fired bursts (Fig. 3A). A reduction of *g*_*BK*_ lead to a broadening of the APs in MEDAKA 1 and MEDAKA 2, similar to what we saw in the experiments (Fig 2C), while it had the opposite effect on the APs fired by RAT. For an intermediate value of *g*_*BK*_, MEDAKA 2 fired broader APs (but still no bursts), MEDAKA 1 became a consistent burster, while only about half of the AP-events in RAT were enduring enough to be classified as bursts (Fig. 3B). When *g*_*BK*_ was set to zero, RAT became consistently non-bursty, while the two medaka models became consistent bursters (Fig. 3C).

**Fig 3.**
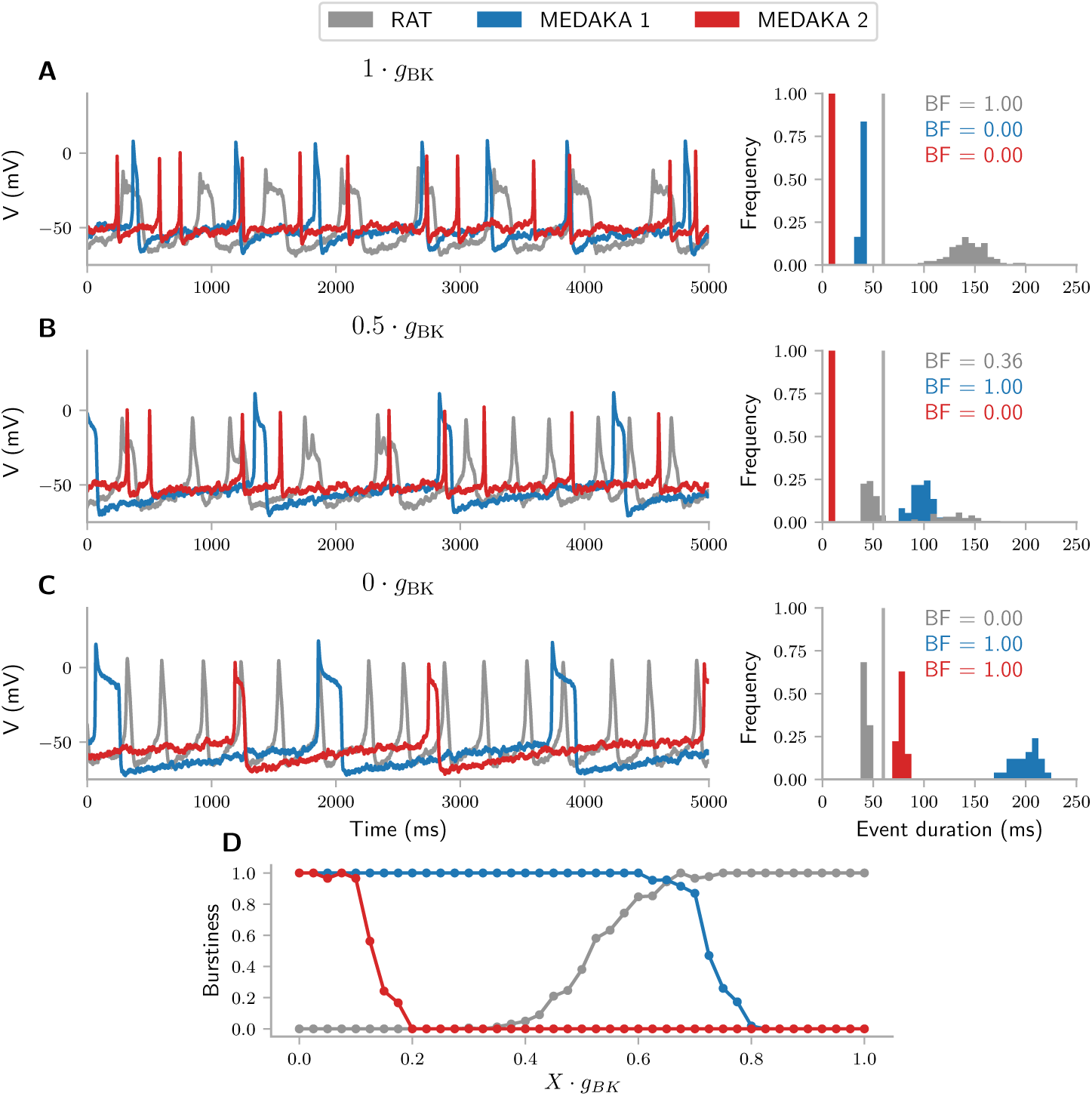
Effects of *I*_*BK*_ on bursting in rat versus medaka. The spontaneous activity of three pituitary cell models, RAT, MEDAKA 1 and MEDAKA 2, for different levels of BK expression. (A) Simulations performed with maximally expressed *g*_*BK*_. (B) Simulations with *g*_*BK*_ reduced to half of the maximum value. (C) Simulations with *g*_*BK*_ set to zero. (A-C) Right panels show histograms of the distribution of AP durations in the respective models. The burstiness factor (BF) denotes the fraction of events that were counted as bursts, i.e., events with a duration exceeding 60 ms. (D) The relationship between *g*_*BK*_ and the propensity for eliciting bursts in the three models. (A-D) The BK conductance is given relative to the maximum value *g*_*BK*_ used in the respective models and was four times bigger in MEDAKA 2 than in RAT and MEDAKA 1. Simulations were otherwise the same as in Fig. 1 of the study by Tabak et al. [9], and a Gaussian noise stimulus was added.

As summarized in Fig. 3D, the relationship between *g*_*BK*_ and the propensity for eliciting bursts in RAT was the opposite of that in the two medaka models. As RAT and MEDAKA 1 differed only in terms of their AP generating mechanisms, these findings suggest that BK has opposing effects on Ca^2+^ generated versus Na^+^ generated APs.

### Why BK currents affect Na^+^ and Ca^2+^ spikes differently

The simulations in Fig. 3 had noise added to them. In order to explore why the same *I*_*BK*_ could have such different effects in RAT versus MEDAKA 1 and MEDAKA 2, we ran the corresponding simulations without noise. In this way, we could ensure that all aspects of the simulated traces reflected membrane mechanisms (and not random fluctuations). In most of the noise-less simulations, all APs within a given spike train were identical (Fig. 4). The models were quite different in terms of their AP firing frequencies, but we here focus predominantly on how BK affected the shape of single APs.

**Fig 4.**
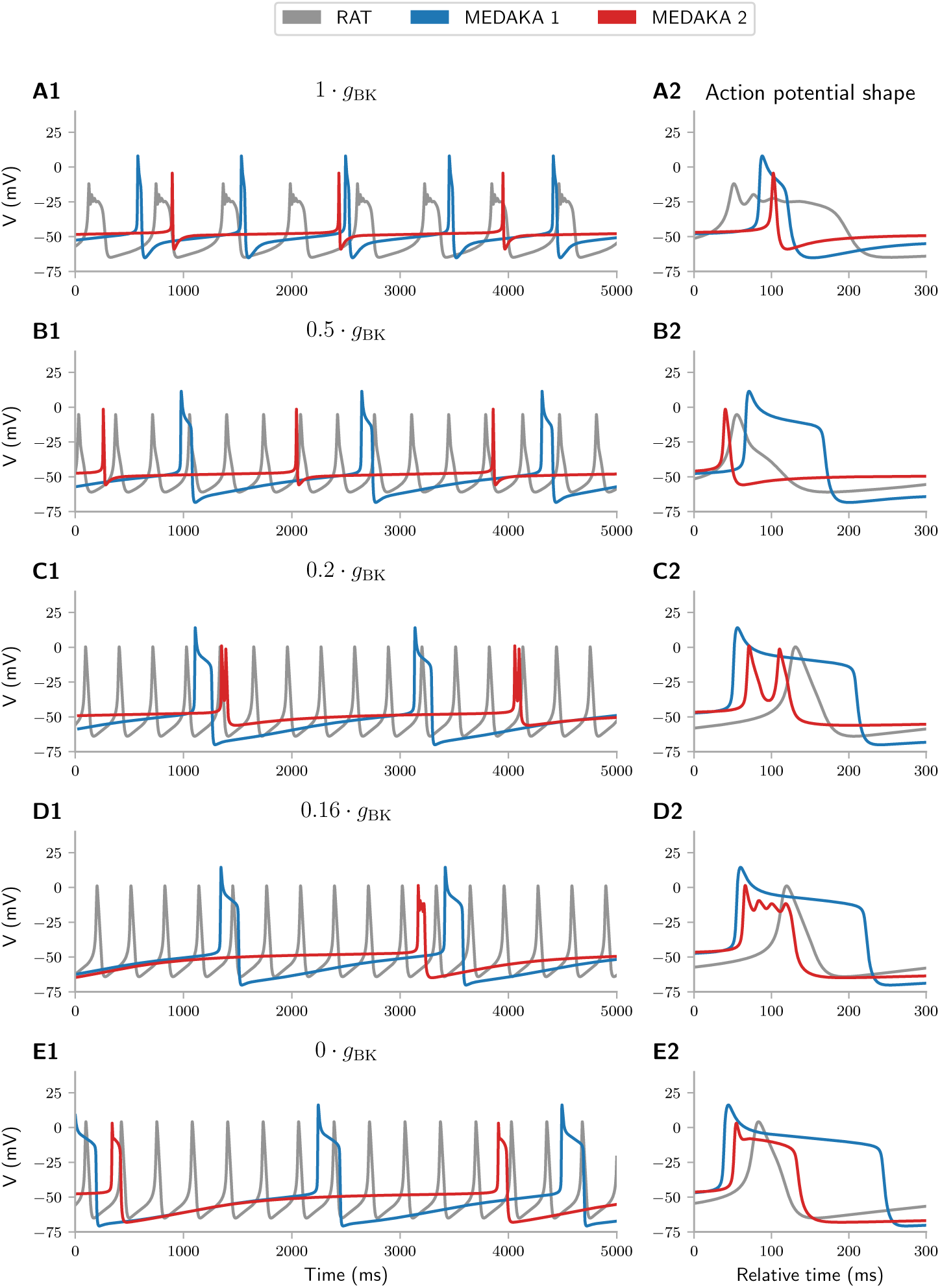
Effects of *I*_*BK*_ of AP shape in rats versus medaka models. (A1-E1) The spontaneous activity of three pituitary cell models, RAT, MEDAKA 1 and MEDAKA 2, for different levels of BK expression (no noise added to the simulations). (A2-E2) Close-ups of selected AP events. The events were aligned, i.e. *t* = 0 (in A2-E2) was defined independently for each trace (in A1-E1) to a point directly before the onset of the selected event. The BK conductance is given relative to the maximum value *g*_*BK*_ used in the respective models and was four times bigger in MEDAKA 2 than in RAT and MEDAKA 1.

The explanation of how BK can act to broaden APs in RAT was given in previous studies [9, 23]. In brief, *I*_*BK*_ acts to reduce the peak amplitude of an AP, as the simulations shown here also demonstrated (compare grey curves in e.g., Fig. 4A2 and E2)). This, in turn, affects *I*_*K*_. Since *I*_*K*_ activates at high voltage levels, the *I*_*BK*_-reduced AP amplitude leads to less *I_K_* activation, and via that to a net reduction in the hyperpolarizing currents active during the AP downstroke. Accordingly, the downstroke occurs more slowly, and the AP event gets broader. In the case with full *g*_*BK*_ expression, the interplay between the depolarizing 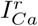 and hyperpolarizing *I*_*BK*_ in RAT also lead to transient oscillations during the downstroke (Fig. 4B2)).

To explain why *I*_*BK*_ acted so differently in the two medaka models, we start by noting that the *I*_*Na*_-mediated upstrokes of their APs were much faster than the 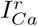-mediated AP in RAT (blue and red curves have faster upstrokes than the grey curves in Fig. 4A2-E2)). *I*_*BK*_, having a time constant of 5 ms in all models, therefore had less time to respond during the rapid AP upstrokes in MEDAKA 1 and MEDAKA 2, and its effect on the AP amplitude was, therefore, smaller than in RAT. Although *I*_*BK*_ did reduce the AP amplitude in all models, the amplitude reduction when going from max *g*_*BK*_ to zero *g*_*BK*_ was by about 16 mV in RAT against only 8 mV and 7 mV in MEDAKA 1 and MEDAKA 2, respectively. Another consequence of the faster AP upstroke in the medaka models was that a major part of the action of *I*_*BK*_ occurred after the AP peak, i.e., during the AP downstroke. In MEDAKA 1 the effect of *I*_*BK*_ was thus shifted away from reducing the AP peak towards contributing to the AP downstroke, and, in this way, *I*_*BK*_ acted to make AP events narrower.

For the case with full *g*_*BK*_ expression (Fig. 4A), MEDAKA 1 fired APs with an unrealistically long duration and at a higher frequency than seen in the experimental data (Fig 1A-B). Even for full BK expression, the AP peaks in MEDAKA 1 were followed by a plateau potential mediated by 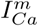 (Fig 4A2). Such *I*_*Ca*_-mediated plateau potentials have been seen in pituitary cells of Atlantic cod [7], but not in goldfish [4] or tilapia [5], and was not seen during spontaneous firing in medaka (Fig. 1A-B). In comparison, MEDAKA 2 had a firing frequency of about 0.7 Hz, with APs peaking at −4.4 mV, and with an AP duration (taken at half max amplitude between −50 mV and AP peak) of 6.4 ms for full BK expression (Fig. 4A). These values were within the range of peak and duration values observed in the experimental data (Fig 1), and as such, MEDAKA 2 was a proper representative for a medaka gonadotrope. MEDAKA 2 also responded to *I*_*BK*_ blockage in a way that resembled that seen in experiments, i.e., *I*_*BK*_ blockage lead to a broadening of the APs (Fig 1D). The resemblance with data was strongest for the simulations with partial blockage, when the AP plateaus underwent oscillations that presumably reflected an interplay between 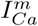 and repolarizing currents activating/inactivating during the downstroke (red lines in Fig 9C2-D2)). It is reasonable to assume that also in the experiments, the blockage of BK by paxilline was not complete. Despite differences between MEDAKA 1 and MEDAKA 2, the effect of *g*_*BK*_ on the AP shape was similar in the two medaka models, and the opposite of that in RAT.

In summary, *I*_*BK*_ had an inhibitory effect on *I*_*K*_ by reducing the AP amplitude, and a collaborative effect with *I*_*K*_ in mediating the AP downstroke. With fast AP upstrokes, as in RAT, the inhibitory effect of *I*_*BK*_ on *I*_*K*_ was stronger than the collaborative, while with slow AP upstrokes, as in MEDAKA 1 and MEDAKA 2, the collaborative effect was stronger than the inhibitory. In this way, *I*_*BK*_ acted as a mechanism that reduced the duration of already brief (Na^+^-mediated) APs, and prolonged the duration of already slow (Ca^2+^-mediated) APs.

### Membrane mechanisms responsible for burstiness

In order to explore in further detail how the various membrane mechanisms affected the AP firing in RAT, MEDAKA 1 and MEDAKA 2, we performed a feature-based sensitivity analysis of the three models. We then assigned the maximum conductances of all included currents uniform distributions within intervals −50% of their default values (Table 1), and quantified the effect that this parameter variability had on selected aspects of the model output (see Methods). An exception was made for *g*_*BK*_, which was assigned a uniform distribution between 0 and the maximum values given in Table 1), i.e., from fully available to fully blocked, motivated by the fact that *g*_*BK*_ did not have a default value in RAT. We note the total order Sobol sensitivity indices considered in the current analysis reflects complex interactions between several nonlinear mechanisms, and that mechanistic interpretations therefore are difficult. Below, we have still attempted to extract the main picture that emerged from the analysis.

Three features of the model responses were considered: (i) *Isbursting*, (ii) *Isregular*, and (iii) *Isfiring*. All these features were binary, meaning e.g., that *Isbursting* was equal to 1 in a given simulation if it contained one or more bursts, and equal to 0 if not. The mean value of a feature (taken over all simulations) then represented the fraction of simulations that had this feature. For example, in RAT *Isbursting* had a mean value of 0.57, *Isregular* had a mean value of 0.37, and *Isfiring* had a mean value of 0.91. This means, respectively, that 57% of the parameterizations of RAT fired bursts, 37 % fired regular APs, and 91% fired APs that were either bursts or regular spikes. We note that the mean values for the *Isbursting* and *isregular* features in RAT and MEDAKA 2 did not sum up exactly to the mean value for the *AP firing* feature. This was because some parameterizations of these two models fired both bursts and regular APs within the same simulation.

AP firing was seen quite robustly in RAT, which fired APs in 91 % of the sampled parameter combinations (mean value of 0.91 in Fig 5A3), while APs were fired in only 77 % and 48% of the parameterizations of MEDAKA 1 and MEDAKA 2, respectively (Fig 5B3-C3). In MEDAKA 2 there was thus AP activity in less than half of the parameterizations. This reflects that the default configuration had a resting potential only slightly above the AP generation threshold, so that any parameter sample that would make the cell slightly less excitable, would abolish its ability for AP generation.

**Fig 5.**
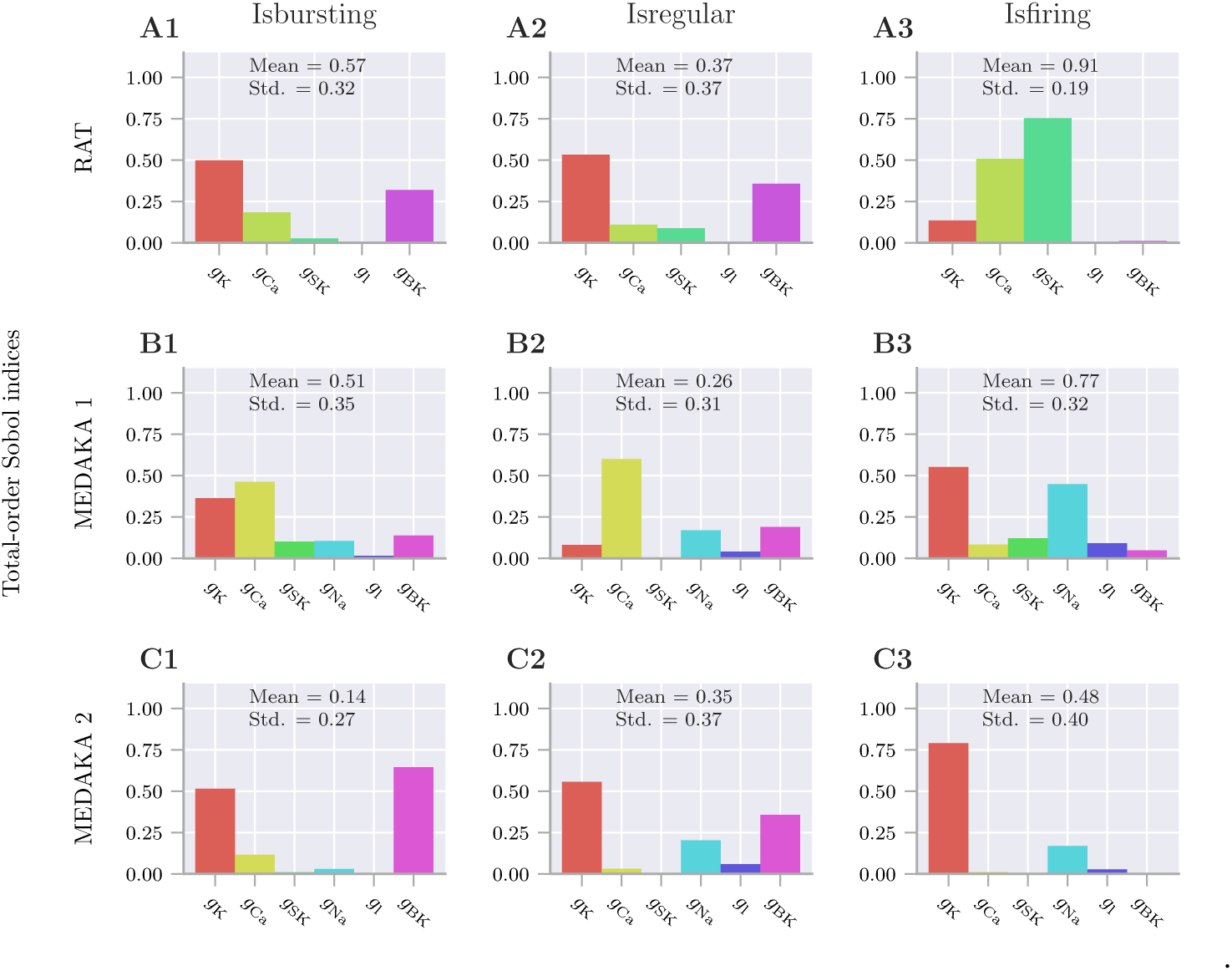
Feature-based sensitivity analysis. Sensitivity of RAT (A), MEDAKA 1 (B) and MEDAKA 2 (C) to variations in the maximum ion channel conductances. The analysis summarizes a large number of simulations where the maximum conductances of all included ion channels were varied within intervals ±50% of their original values. An exception was made for *g*_*BK*_, which was varied between 0 and 0.32 mS/cm^2^ in (A) RAT and (B) MEDAKA 1, and between 0 and 1.26 mS/cm^2^ in (C) MEDAKA 2. (A1-C1) *Isbursting* denotes the fraction of simulations that elicited one or more bursts, (A2-C2) *Isregular* denotes the fraction of simulations that elicited one or more regular APs, and (A3-C3) *Isfiring* denotes the fraction of simulations that elicited any AP events at all. (A-C) Histograms depict the total order Sobol sensitivity indices (see Methods). The analysis was performed by aid of the recently developed toolbox *Uncertainpy* [34] (see Methods for details). Simulations were run with no noise.

Within the explored parameter range, RAT and MEDAKA 1 tended to be bursting, meaning that bursts were seen in a larger fraction of the simulations than regular spikes (mean values in Fig 5A1 and B1 are larger than in Fig 5A2 and B2). MEDAKA 2, on the contrary, was adapted to experimental data where AP firing under control conditions (Fig. 1A-B) was regular and with very narrow APs. MEDAKA 2 thus had a high propensity for firing regular APs and elicited bursts only in (14%) of the parameter combination (Fig 5C1-C2).

The total order Sobol indices (shown in histograms in Fig 5) quantify how much of the variability (between different simulations) in the response features that were explained by the variation of the different model parameters (i.e., maximum conductances). For example, *Isfiring* in MEDAKA 1 and MEDAKA 2 were most sensitive to *g_K_* and *g_Na_* (Fig 5B3 and C3)), meaning that these conductances were most important for whether the model was capable of generating AP events. This result was not surprising, since *I*_*K*_, *I*_*Na*_ (along with the leakage current, *I_l_*) were the only currents with a nonzero activity level around rest, and thus were the ones that determined whether the resting potential was above the AP firing threshold (cf. Fig 2).

Having other AP generation mechanisms, *Isfiring* in RAT was instead sensitive predominantly to 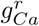 and *g*_*SK*_ (Fig 5A3)). Due to the wide activation range of 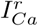, generating the APs in RAT, this model was in principle always above the AP firing threshold. The instances when AP firing was not seen in RAT then reflected lack of hyperpolarizing mechanisms, such as *g*_*SK*_, that could repolarize the membrane potential after the AP peak.

In general, *Isbursting* and *Isregular* were not sensitive to the same parameters as *Isfiring*. This implies that the mechanisms determining whether the cell fired an AP or not were not the same as those determining the duration of the AP once it had been fired. For example, *Isfiring* in MEDAKA 2 was nearly insensitive to *g*_*BK*_, while *Isbursting* and *Isregular spiking* were highly sensitive to this parameter. A little simplified, we may say that *g*_*K*_ and *g*_*Na*_ thus determined whether MEDAKA 2 fired an AP (Fig 5C3), while *g*_*BK*_ (and to some degree 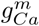) determined whether the AP became a burst or a regular spike (Fig 5C1-C2). In MEDAKA 1, the situation was more complex, and its *Isbursting* and *Isregular* were sensitive to almost all model parameters (Fig 5B1-B2).

In RAT, *Isbursting* was most sensitive to *g*_*K*_ and second most to *g*_*BK*_ (Fig 5A1). The lower sensitivity to *g*_*BK*_ compared to *g*_*K*_ might seem surprising, given the previously proclaimed role of *g*_*BK*_ as a burst promoter in this model [9]. However, we can interpret this finding in light of the above-established understanding of burst generation in RAT. We remember that *I*_*BK*_ promoted bursting indirectly by inhibiting *I*_*K*_, so that final determinant for whether an event became a burst was the magnitude (or rather the reduced magnitude) of *I*_*K*_. The results from the sensitivity analysis then simply indicate the direct effect on *I_K_* obtained by a variation of *g*_*K*_ was larger than the indirect effect on *I*_*K*_ obtained by variation of *g*_*BK*_.

We note that the three different feature sensitivities in Fig 5 were not independent. For example, if variations in a given parameter tended to switch the cell between bursting and not firing at all, *Isbursting* and *Isfiring* would share a high sensitivity to this parameter. Likewise, if variations in a given parameter instead tended to switch the cell between bursting and regularly spiking, *Isbursting* and *Isregular* would share a high sensitivity to this parameter. In this regard, *g*_*BK*_ was the parameter with the cleanest role as a switch between bursting and regular spiking, as in all models, the *Isbursting* and *Isregular* had a high sensitivity to *g*_*BK*_, while *Isfiring* had a quite low sensitivity to *g*_*BK*_.

A more general insight from the sensitivity analysis was that the propensity for AP firing, regular spiking and bursting in all three models depended on a complex interplay between several mechanisms. All models could be shifted from regularly spiking to bursty by changes, not only in *g*_*BK*_ (as demonstrated in Fig 3 and Fig 4), but also in other conductances (such as *g*_*K*_ or *g_Ca_*). In all models, however, bursts were facilitated by *g_Ca_* and counteracted by *g*_*K*_, while *g*_*BK*_ was the more fascinating mechanism, having the opposite effect on the burstiness in RAT versus the two medaka models.

## Discussion

TTX-sensitive Na^+^ currents (*I*_*Na*_) are present in all pituitary cells, but are in many cases inactive during spontaneous activity [8]. Previous models of the electrical activity of pituitary cells have focused on conditions where *I*_*Na*_ is of lesser importance, and where AP generation is predominantly mediated by high-voltage activated Ca^2+^ currents [3, 9, 22–26]. To our knowledge, we have in the current work presented the first models that describe pituitary cells under conditions where AP generation is *I*_*Na*_-mediated. The model was adapted to experimental data from gonadotrope luteinizing hormone-producing cells in medaka, whose spontaneous activity is highly *I*_*Na*_-dependent. Voltage-clamp data was used to develop models for the activation kinetics for *I*_*Na*_ and *I*_*Ca*_ currents, and the firing properties of the model were further adapted to current-clamp data from spontaneously active cells (under control conditions, and after application of TTX and paxilline).

An important goal of this work was to perform a comparison between the response properties of pituitary cells that elicited *I*_*Ca*_-mediated APs and those that elicited *I*_*Na*_-mediated APs. For this, we used a model based on data from rat, which elicited *I*_*Ca*_-mediated APs, and compared it to two models (MEDAKA 1 and MEDAKA 2) based on data from medaka, both of which elicited *I*_*Na*_. mediated APs. In this context, MEDAKA 1 had the (methodological) advantage that all hyperpolarizing membrane mechanisms were kept identical to those in RAT, so that RAT and MEDAKA 1 differed only in terms of AP generation mechanisms. MEDAKA 2 was more strongly adapted to experimental data, and had the advantage that its firing properties were more representative for real medaka gonadotropes. The most interesting result that came out of this comparison, was that BK currents (*I*_*BK*_) had a diametrically opposite role in terms of how they affect the AP shape in RAT versus the medaka models. When APs were generated by *I*_*Ca*_ (as in RAT), they were relatively slow, and *I*_*BK*_ then acted to broaden the AP events and promote bursting behavior [9, 23]. On the contrary, when APs predominantly were generated by *I*_*Na*_ (as in MEDAKA 1 and MEDAKA 2), they had a rapid upstroke, and *I*_*BK*_ then acted to make the AP narrower, and prevent bursting behavior (Fig 3). This suggests that *I*_*BK*_ acts as a mechanism that distinguishes between rapid and slow APs, and amplifies the difference between the two by narrowing down already narrow APs while broadening already broad APs.

It should be noted that the effect seen in medaka gonadotropes, i.e., that *I*_*BK*_ acted to reduce AP duration, is a commonly reported role for *I*_*BK*_ in many excitable cells [35–39], while the burst promoting effect that *I*_*BK*_ had in rat pituitary cells [9, 23] is less conventional. Furthermore, the role of *I*_*BK*_ as a burst promoter has not been found consistently in rat pituitary cells. In the study by Miranda et al. 2003, AP duration in rat pituitary cells was instead found to increase when *I*_*BK*_ was blocked with paxilline [38], i.e. similar to what we found for medaka gonadotropes (Fig 1D). The different effects of *I*_*BK*_ on spike duration observed in different laboratories [23, 35, 38] was addressed by Tabak et al. 2011 [9], who proposed possible explanations that could reconcile the conflicting results. One possible explanation could be there is a variability in terms of how BK channels are localized in various cells, and that BK channels that are co-localized with Ca^2+^ channels will respond rapidly to voltage fluctuations and promote bursting, while BK channels that are not co-localized with Ca^2+^ channels will react more slowly to voltage fluctuations and have the opposite effect [9]. A second possible explanation, also suggested by Tabak et al. 2011, was that *I*_*BK*_ might have different kinetic properties in different cells due to variations in their phosphorylation state [9]. A third explanation could be that different cells have different BK splice variants [40], or different regulatory sub-units.

The simulations presented in Fig 3 provide an alternative possible explanation to the conflicting conclusions regarding the role of *I*_*BK*_. The fact that *I*_*BK*_ has a different effect on the AP shape in different cells does not by necessity reflect a variation in *I*_*BK*_ expression or kinetics between the different cell types, as proposed by Tabak et al. 2011. As Fig 3 showed, the same model of *I*_*BK*_ could either prolong or reduce AP generation, depending on which membrane mechanisms (*I*_*Ca*_ or *I*_*Na*_) that mediated the AP upstroke. Such differences in AP mechanisms could in principle explain the differences between the conflicting experiments on rat pituitary cells [23, 38]. In the experiment by Van Goor et al. 2001, where *I*_*BK*_ was found to prolong the AP duration, APs were predominantly mediated by *I*_*Ca*_ [9, 23]. In the experiment by Miranda et al. 2003, where *I*_*BK*_ was found to shorten the AP duration (i.e., that blocking *I*_*BK*_ lead to longer APs), it was reported that this *only* occurred under conditions in which short APs were present. It is likely that the events described in that work as *short APs*, were *I*_*Na*_-mediated APs, so that the differences between the two studies by Van Goor 2011 and Miranda 2003 are reflected by the differences between the simulations on RAT versus MEDAKA 1 and MEDAKA 2 in Fig 3.

Although MEDAKA 2 captured the essential firing properties of medaka gonadotropes, the agreement between model and data was not perfect. For example, the AP duration in MEDAKA 2 during control conditions (Fig 9C) was in the upper range of that seen in the experiments (Fig 1A-B), and we were not able to obtain briefer APs in the model without compromising the agreement between the experimental data and other model features, such as the AP peak amplitude, afterhyperpolarization, and response to *I*_*BK*_-blockage. The conductances selected for MEDAKA 2 (Table 1) were thus a compromise made to obtain an acceptable match to several features simultaneously. The fact that we were not able to obtain a more accurate match between model and data likely reflect that some of the ion channels present in the model are imperfect representations of the ion channels present in the real cell. For example, the simplified kinetics schemes used for *I*_*K*_, *I*_*BK*_ and *I*_*SK*_ were adopted from a model of rat pituitary cells [9], and were not constrained to data from medaka gonadotropes. In addition, the biological cell is likely to contain a variety of additional ion channels [8] that were not included in the model.

To our knowledge, MEDAKA 1 and MEDAKA 2 are the first computational models that describe *I*_*Na*_-based APs in endocrine cells. Although MEDAKA 2 was adapted to experimental recordings from gonadotrope luteinizing hormone-producing cells in medaka, we believe that the model can have a more general value. Different types of pituitary cells in several different species share many of the same membrane mechanisms [8]. In particular, *I*_*Na*_-based APs are elicited by several pituitary endocrine cell types and in several animal species, depending on biological conditions [4, 7, 17, 18, 30–32]. It is thus likely that the response patterns of related cell types may be captured by up- or down-regulation of selected mechanisms included in MEDAKA 2. In that context, the sensitivity analysis presented in Fig 5 may give useful guidance in terms of which parameters that should be adjusted in order to obtain desired changes in the models firing properties.

## Methods

### Experimental procedures

The electrophysiological experiments were conducted using the patch-clamp technique on brain-pituitary slices from adult female medaka (as described in [41]). To record spontaneous action potentials and Ca^2+^ currents we used amphotericin B perforated patch configuration, while for Na^+^ currents we used whole cell configuration. Extracellular (EC) solution used for recording spontaneous action potentials (current clamp) contained 134 mM NaCl, 2.9 mM KCl, 2 mM CaCl_2_, 1.2 mM MgCl_2_ 1.2, 10 mM HEPES, 4.5 mM glucose. The solution was adjusted to a pH of 7.75 with NaOH (added drop-wise from 1M solution) and osmolality adjusted to 290 mOsm with mannitol before sterile filtration. Before use, the EC solution was added 0.1 % bovine serum albumin (BSA). For Na^+^ current recordings (voltage clamp) we used a Ca^2+^ free and Na^+^ fixed (140 mM) EC solution, pH adjusted with trizma base. In addition, 10 *µ*M nifedipine, 2 mM 4-Aminopyridine (4-AP) and 4 mM Tetraethylammonium (TEA) was added to the EC solution just before the experiments. To record Ca^2+^ currents, we substituted NaCl with 120 mM choline-Cl and added 20 mM Ca^2+^, 2 mM 4-AP and 4 mM TEA. The patch pipettes were made from thick-walled borosilicate glass using a horizontal puller (P 100 from Sutter Instruments). The resistance of the patch pipettes was 4-5 MΩ for perforated patch recordings and 6-7 MΩ for whole-cell recordings. For recordings of spontaneous action potentials, the following intracellular (IC) electrode solution was added to the patch pipette: 120 mM KOH, 20 mM KCl, 10 mM HEPES, 20 mM Sucrose, and 0.2 mM EGTA. The pH was adjusted to 7.2 using C^6^H^13^NO^4^S (mes) acid, and the osmolality to 280 mOsm using sucrose. The solution was added 0.24 mg/ml amphotericin B to perforate the cell membrane (see [41] for details). In voltage clamp experiments the K^+^ was removed from the intracellular solution to isolate Na^+^ and Ca^2+^ currents. This was achieved by substituting KOH and KCl with 130 mM Cs-mes titrated to pH 7.2 with CsOH. The electrode was coupled to a Multiclamp 700B amplifier (Molecular Devices) and recorded signal was digitized (Digidata 1550 with humsilencer, Molecular Devices) at 10 KHz and filtered at one-third of the sampling rate. In selected experiments, voltage-gated Na^+^ channels were blocked using 5 *µ*M TTX, and BK channels were blocked using 5 *µ*M paxilline. Both drugs were dissolved in EC solution and applied using 20 kPa puff ejection through a 2 MΩ pipette, 30-40 *µ*m from the target cell.

Under the experimental (voltage-clamp) conditions used for recording Na^+^ currents, and under the experimental current-clamp conditions, a liquid junction potential of about *−*9 mV was calculated and corrected for in the data shown in Fig 1, and in the kinetics model for *I*_*Na*_ (Fig 2A). A liquid junction potential of about −15 mV was calculated for the experimental (voltage-clamp) conditions used for recording Ca^2+^ currents, and was corrected for in the kinetics model for 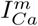 (Fig 2B).

### Models

As stated in the Results-section, RAT was described by the equation:

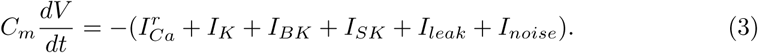

and the two medaka models, MEDAKA 1 and MEDAKA 2, by:

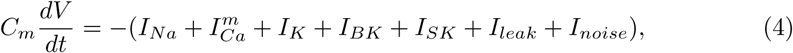

The passive membrane properties were the same in all models, with specific membrane capacitance *C_m_* = 3.2 *µ*F*/*cm^2^, and a leak conductance 0.032 mS*/*cm^2^.

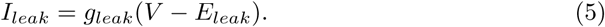

RAT had a passive reversal potential of *E_leak_* = *−*50 mV. In RAT, 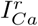 was quite active at the resting potential, and counteracted the hyperpolarizing effect of *I*_*K*_ which was also had a non-zero activity level around the resting potential. Replacing 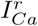 with 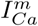 in MEDAKA 1 and MEDAKA 2 therefore removed a source of spontaneous depolarization and lead to an unwanted hyperpolarization of the resting potential. To compensate for this, *E_leak_* was adjusted to −45 mV in MEDAKA 1 and MEDAKA 2, which resulted in an effective resting potential around −50 mV, similar to that in RAT, and also to the experimental data in Fig 1.

Below, we give a detailed description of the active ion channels. The kinetics of all ion channels were summarized in Fig. 2, the model parameters that differed between the models were listed in Table 1, while new parameters will be defined when introduced.

*I*_*Na*_ (MEDAKA 1 and MEDAKA 2) was modeled using the standard Hodgkin and Huxley-form [42]:

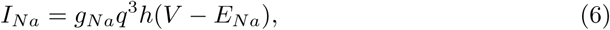

with a reversal potential *E_Na_* = 50 mV, and gating kinetics defined by:

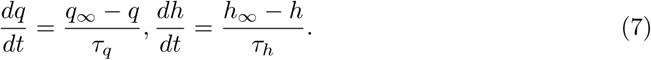

The steady-state activation and time constants (*q_∞_*, *h_∞_*, *τ_q_* and *τ_h_*) were fitted to voltage-clamp data from medaka gonadotropes, as described below, in the subsection titled “Model for the voltage-gated Na^+^ channels”.

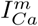 (MEDAKA 1 and MEDAKA 2) was modelled using the Goldman-Hodgkin-Katz formalism, which accounts for dynamics effect on Ca^2+^ reversal potentials [43]:

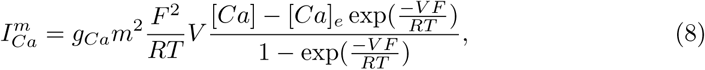

with

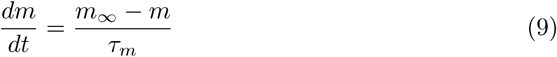

Here, *R* = 8.314J*/*(mol · K is the gas constant, *F* = 96485.3C*/*mol is the Faraday constant, *T* is the temperature, which was set to 293.15 K in all simulations. [*Ca*] and [*Ca*]_*e*_ were the cytosolic and extracellular Ca^2+^ concentrations, respectively. The former was explicitly modelled (see below), while the latter was assumed to be constant at 2 mM. As Eq 8 shows, we used two activation variables *m*. This is typical for models of L-type Ca^2+^ channels (see e.g. [44–47]), which are the most abundantly expressed HVA channels in the cells studied here [11]. The steady-state activation and time constant (*m_∞_* and *τ_h_*) were fitted to voltage-clamp data from medaka gonadotropes, as described below, in the subsection titled “Model for high-voltage activated Ca^2+^ channels”. We note that *g_Ca_* in the Goldman-Hodgkin-Katz formalism (Eq 8) is not a conductance, but a permeability with units cm/s. (It is proportional to the conductance, and for simplicity, we have referred to it as a conductance in the text).

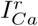 (RAT) was a simpler Ca^2+^-channel model, with one gating variable and constant reversal potential [9]:

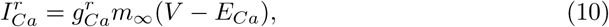

with instantaneous steady-state activation:

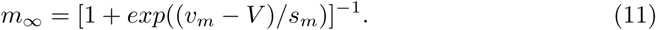

where *s*_*m*_ = is a slope parameter and *v*_*m*_ the potential at half-activation.

The directly rectifying K^+^ channel (all models) was modelled as

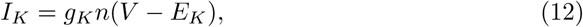

with reversal potential *E_K_* = − 75 mV, and a time dependent activation variable described by

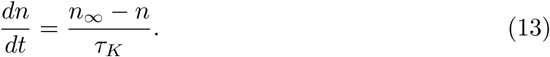

The constant (voltage-independent) time constant *τ*_*K*_ was 30 ms in RAT and MEDAKA 1, and 5 ms in MEDAKA 2, and the steady-state activation was described by:

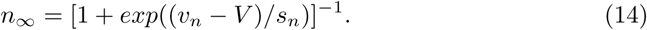

with a slope parameter *s*_*n*_ = 10 mV, and half-activation *v_n_* = −5 mV.

The BK-channel (all models) was modelled as

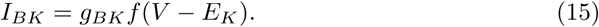

The activation kinetics was:

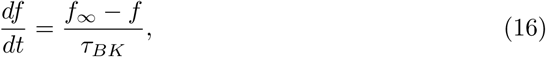

with a constant (voltage-independent) activation time constant *τ_BK_* = 5 ms. The steady-state activation was given by:

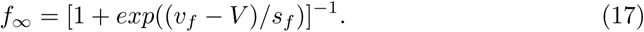

with a slope parameter *s*_*m*_ = 12 mV, and half-activation *v_m_* with values −20 mV in RAT and MEDAKA 1 and −15 mV in MEDAKA 2. We note that BK was modeled as a voltage (and not Ca^2+^) dependent current. The rationale behind this simplification was that BK activation depends on the Ca^2+^ concentration in highly localized Ca^2+^ nanodomains, where it is co-localized with, and depends on influx through, high-voltage activated Ca^2+^ channels. This influx is in turn is largely determined by the membrane potential and reaches an equilibrium within microseconds [9], making the BK channel effectively voltage dependent.

Finally, the SK channel (all models) was modeled as:

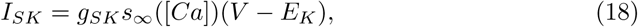

with an instantaneous, Ca^2+^ dependent, steady-state activation:

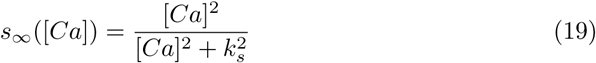

where [*Ca*] denotes the cytosolic Ca^2+^ concentration, and *k*_*s*_ was a half-activation concentration of 0.4 *µ*M.

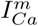 and *I*_*SK*_ were dependent on the global cytosolic Ca^2+^ concentration. This was modelled as a simple extrusion mechanism, receiving a source through 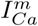 (for MEDAKA 1 and MEDAKA 2) and 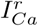 for RAT, and with a concentration dependent decay term assumed to capture the effects of various extrusion and buffering mechanisms:

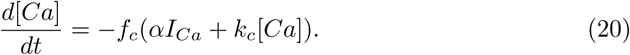

Here, *f_c_* = 0.01 is the assumed fraction of free Ca^2+^ in the cytoplasm, *α* = 0.004725 mM · cm^2^/*µ*C converts an incoming current to a molar concentration, and *k*_*c*_ = 0.12 ms^−1^ is the extrusion rate [9]. Due to the requirements from the NEURON simulator, the parameters from the original model by Tabak et al. [9] were converted from total currents/capacitance to currents/capacitance per membrane area, using a cell body with a membrane area of *π* · 10^−6^ cm^2^ (see https://github.com/ReScience/ReScience-submission/pull/53).

In the simulations in Fig 3, a noise input was added, described by

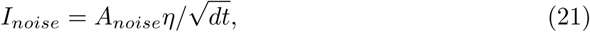

where the noise amplitude (*A*_*noise*_) was 4 pA, and *η* was a random process drawn from a normal distribution [9].

### Model for the voltage-gated Na^+^ channels

The steady-state values and time courses of the gating kinetics were determined using standard procedures (see e.g. [32, 42, 48, 49]), and was based on the experiments summarized in Fig 6. To determine activation, the cell was held at −60 mV for an endured period, and then stepped to different holding potentials between −80 to 100 mV with 5 mV increments (Fig. 6A2), each for which the response current (*I*_*Na*_) was recorded (Fig. 6A1). The inactivation properties of Na^+^ were investigated using stepwise pre-pulses (for 500 ms) between −90 and 55 mV with 5 mV increments before recording the current at −10 mV (Fig. 6B2). The resulting Na^+^ current then depended on the original holding potential (Fig. 6B1). Finally, the recovery time for the Na^+^ current was explored by exposing the cell to a pair of square pulses (stepping from a holding potential of −60 mV to −10 mV for 10 ms) separated by a time interval Δ*t* (Fig. 6C2). The smallest Δ*t* was 10 ms, and after this, Δ*t* was increased with 100 ms in each trial. The cell responded to both pulses by eliciting Na^+^-current spikes (Fig. 6C2). When Δ*t* was small, the peak voltage of the secondary spike was significantly reduced compared to the first spike, and a full recovery required a Δ*t* in the order of 1/2-1 s.

**Fig 6.**
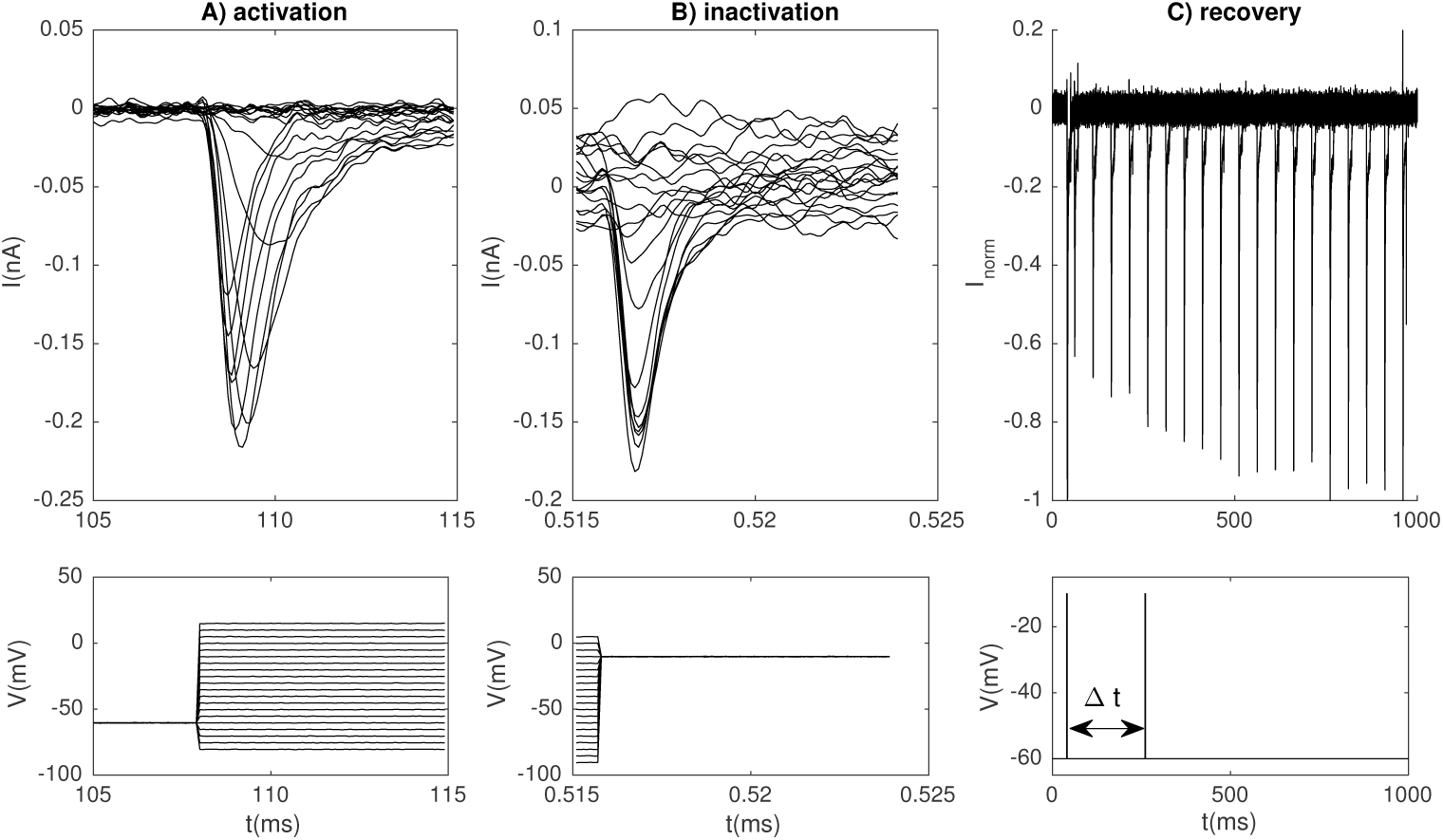
Experimentally recorded Na^+^ currents. (A) Na^+^ currents evoked by the activation protocol. (B) Na^+^ currents evoked by the inactivation protocol. (C) Na^+^ currents used to determine the time constant for recovery from inactivation. (A-C) Voltage protocols are shown below the recorded currents, and all panels show a series of experiments (traces). In (C), the cell was exposed to a pair of square 10 ms pulses arriving with various inter-pulse intervals (Δ*t*). The first pulse always arrived after 40 ms (and coincided in all experiments), while each secondary spike represents a specific experiment (i.e., a specific Δ*t*). The traces were normalized so that the first spike had a peak value of −1 (corresponding to approximately −0.25 pA).

#### Steady-state activation and inactivation

To determine steady-state activation, the peak current (*I*_*max*_) was determined for each holding potential in Fig. 6A, and the maximum peak was observed at about −10 mV. For inactivation, the peak current (*I*_*max*_) was recorded for each holding potential in Fig. 6B. In both cases, the maximal conductance (*g_max_*) for each holding potential was computed by the equation:

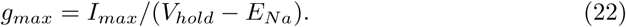

Under the experimental conditions, the intra- and extracellular Na^+^ concentrations were 4 mM and 140 mM, respectively, and the temperature was 26 degrees Celsius, which gives a reversal potential (*E*_*Na*_ = *RT /F ·* ln([Na]_ex_)*/* ln([Na]_in_) of 92 mV. The estimates of *g*_*max*_ for activation and inactivation are indicated by the markers ‘x’ in Fig. 7A and B, respectively, and markers ‘o’ indicate a secondary experiment. The dependency of *g*_*max*_ on *V*_*hold*_ was fitted by a Boltzmann curve:

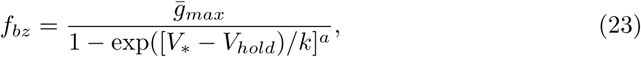

where 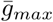 corresponds to *g*_*max*_ estimated for the largest peak in the entire data set (i.e. at about −0.22 nA in Fig. 6A and −0.18 nA in Fig. 6B). The factor *k* determines the slope of the Boltzman curve, the exponent *a* corresponds to the number of activation or inactivation gates, and *V_∗_* determines the voltage range where the curve rises. When *a* = 1 (as for inactivation), *V_*_* equals *V*_1/2_, i.e. the voltage where *f*_*bz*_ has reached its half maximum value. With a higher number of gates, *V*_*_ = *V*_1/2_ + *k* · ln(2^1/*a*^ − 1). Eq 23 gave a good fit for the steady state activation 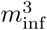 (Fig 7A) and the steady state inactivation *h*_inf_ (Fig 7B) with the parameter values for *a*, *k* and *V*_*_ listed in Table 2.

**Fig 7.**
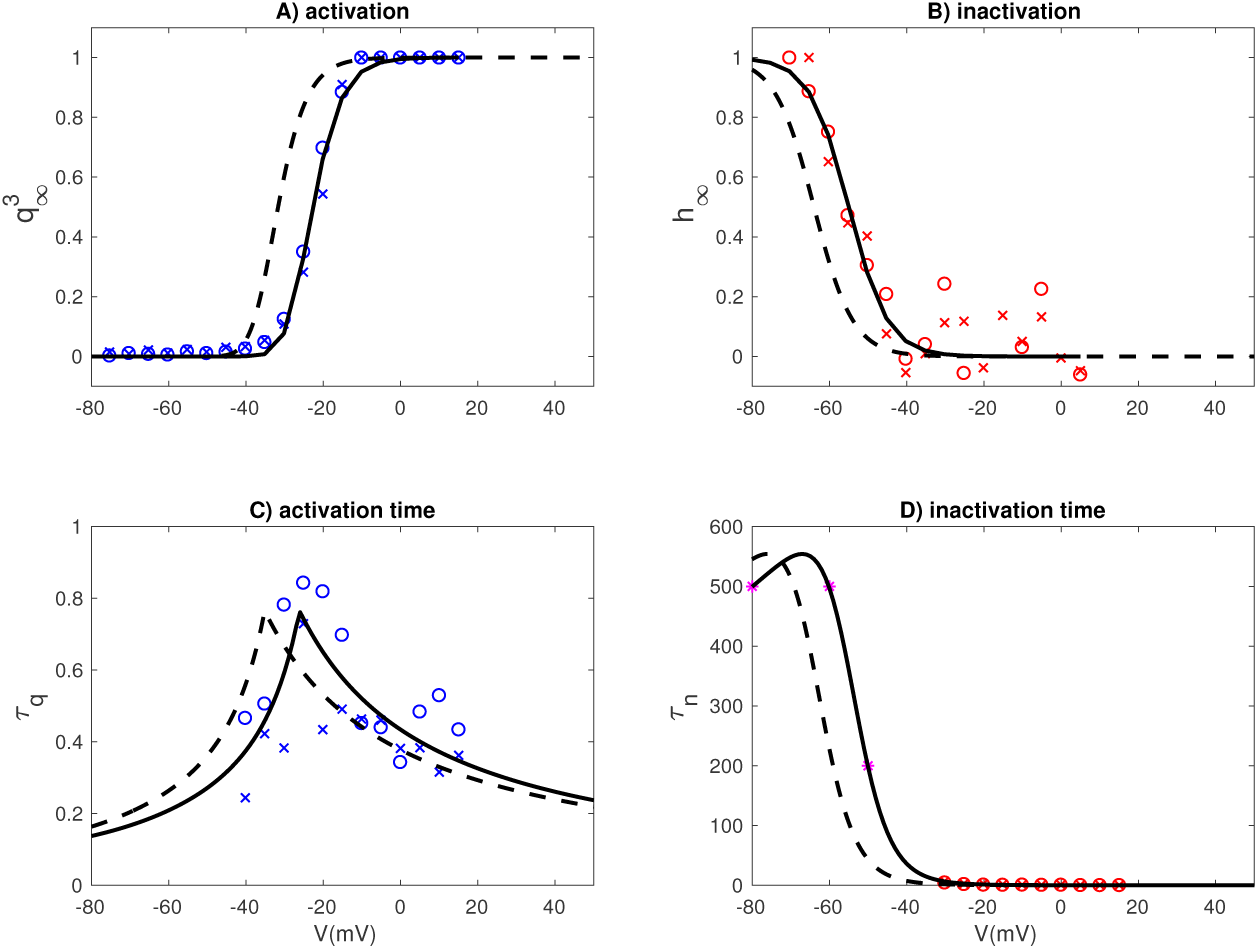
Fitted kinetics for the Na^+^ current. Voltage dependence of steady-state activation (A), steady-state inactivation (B), activation time constant (C) and inactivation time constant (D). The data points and curves in (A-B) were normalized so that activation/inactivation curves had a maximum value of 1 (assuming fully open channels). Dashed lines represent the same model when corrected for a liquid junction potential of 9 mV.

**Table 2.**
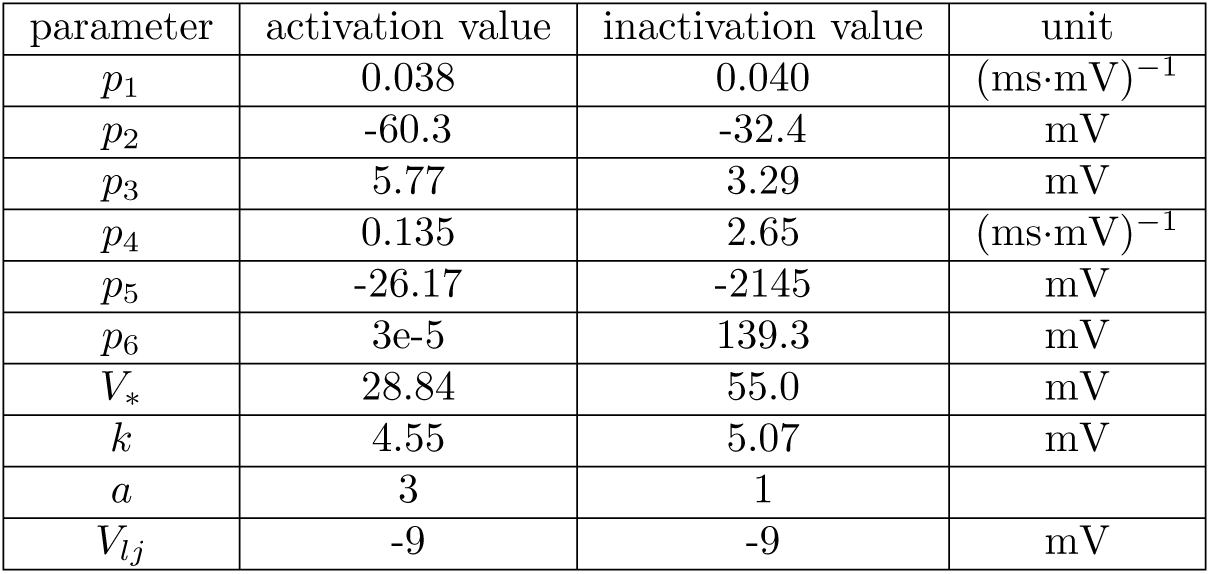
Parameters for Na^+^ activation. The parameters *p*1-*p*6 are used together with Eq. 25 to yield the time constants for steady state activation and inactivation (in units of ms). The remaining parameters are used together with Eq. 23 to obtain the steady-state activation and inactivation functions. The curves obtained in this way describe the voltage dependence under experimental conditions, and was afterwards corrected by subtracting the liquid junction potential of −9 mV (see Fig 2A).

#### Time constants for activation and inactivation

With three opening gates (*q*) and one closing gate (*h*), the time constants for activation and inactivation were derived by fitting the function [42]

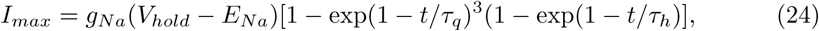

to the response curves in Fig. 6A1. For low step potentials (< −40 mV), the response was too small and noisy to reveal any clear trend, and we were unable to obtain meaningful fits using the functional form of Eq. 22. For this reason, only the experiments with a step potential of −40 mV and higher were used when fitting the time constants. The fitting procedure resulted in a pair of time constants (*τ*_*q*_ and *τ*_*h*_) for each step potential in the protocol, as indicated by the data points (’x’ and ‘o’) in Fig. 7C and D. The data points obtained by fitting Eq. 24 to the traces in Fig. 6A1 were sufficient to obtain a clear picture of the voltage dependence of the activation time constant (*τ*_*q*_), which had a peak value at −24 mV, i.e. within the voltage-range for which there was suitable data (Fig. 7C). The inactivation time constant (*τ*_*h*_) was, however, monotonously decreasing over the voltage range for which there was good data. We therefore needed additional data points for the voltage dependence of *τ*_*h*_ in the range *V* < −40 mV. Based on the insight from the recovery-experiments (Fig. 6C), we expected inactivation to be very slow at the resting potential and below. To account for this, we introduced the three additional data points marked by ‘*’ in Fig. 7D, which assure a recovery time in the correct order of magnitude.

The data points for the time constants were fitted with curves on the functional form proposed by Traub et al. [50]:

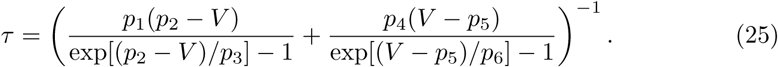

Good fits to the data points were obtained with the parameter values in Table 2.

### Model for high-voltage activated Ca^2+^ channels

When estimating the steady-state values and time constant we followed procedures inspired from previous studies of L-type Ca^2+^ channel activation, we did not use Eq 8, but used the simpler kinetics scheme *I*_*Ca*_ = *g*_*HV A*_*m*^2^(*V – E*_*Ca*_) (see e.g. [45])) assuming a constant reversal potential.

#### Steady-state activation

The steady-state value and time constant for *m* were determined from the experiments summarized in Fig. 8A-B. To study steady-state activation, the cell was held at −60 mV for an endured period, and then stepped to different holding potentials, each for which the response current (*I*_*Ca*_) was recorded (Fig. 8A). Due to the small cellular size, perforated patches was used for recording the Ca^2+^ currents, and the recorded currents were small and noisy. As Fig. 8A shows, the *I*_*Ca*_ responses did not follow a characteristic exponential curve towards steady state, as seen in many other experiments. Likely, this was due to *I*_*Ca*_ comprising a complex of different HVA channels (e.g., P, Q, R, L-type) which have different activation kinetics [44–46, 51–53]. In addition, some in some of the weaker responses *I*_*Ca*_ even switched from an inward to an outward current, something that could indicate effects of ER release on the calcium reversal potential. Due to these complications, only the early part of the response was used, i.e., from stimulus onset and to the negative peak value in interval indicated by dashed vertical lines). Voltage-dependent deactivation of Ca^2+^ currents (Fig. 8B) was examined by measuring the tail current that followed after a 5 ms step to 10 mV when returning to voltages between −10 mV and −60 mV. The deactivation protocol was used to provide additional data points for the activation time constants (see below).

**Fig 8.**
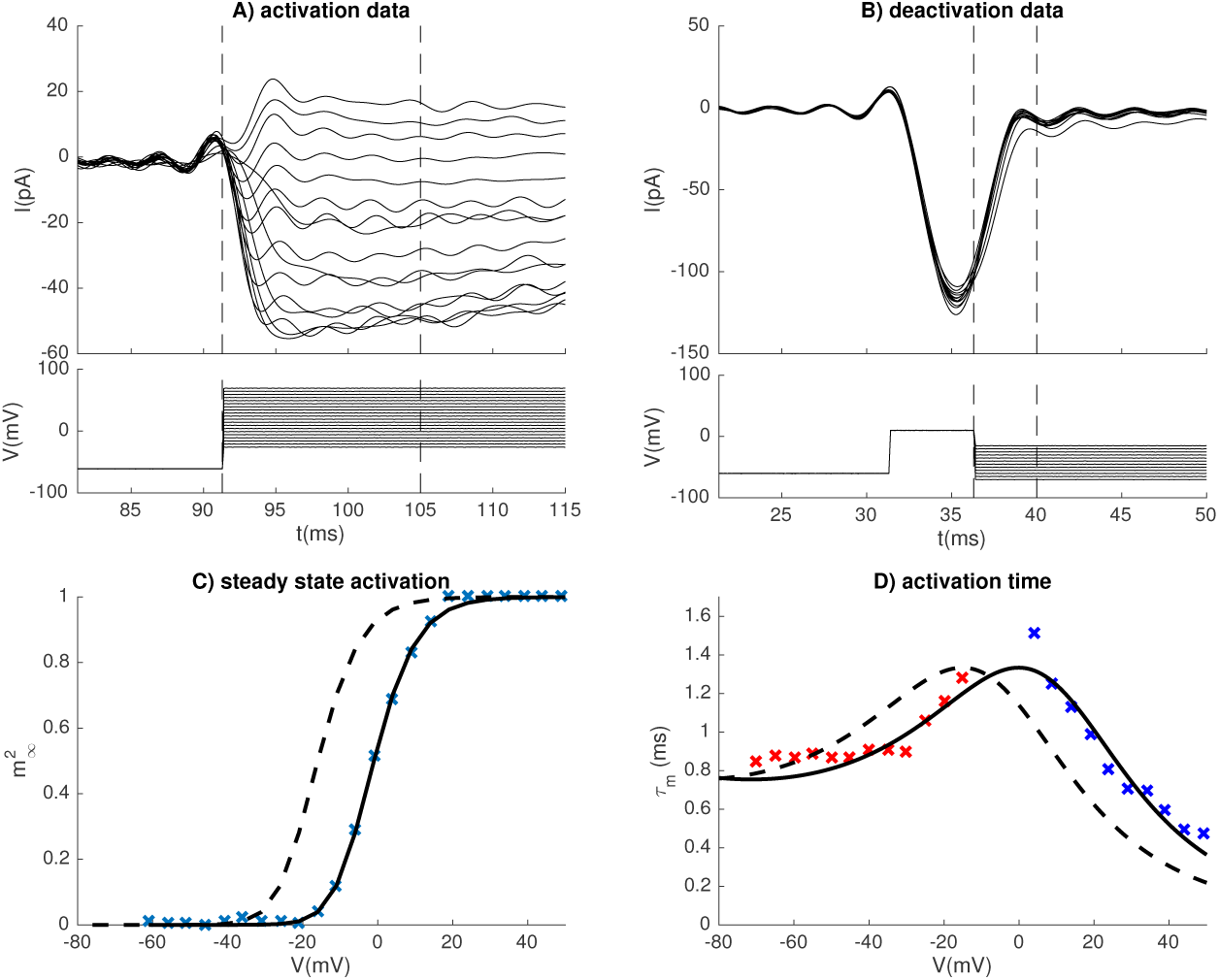
Fitted kinetics for the Ca^2+^ current. (A) *I*_*Ca*_ evoked by the activation protocol. (B) *I*_*Ca*_ evoked by the deactivation protocol. (A-B) Voltage protocols are shown below the recorded currents, and all panels show a series of experiments (traces). The current-traces were low-pass filtered with a cutoff-frequency of 300 Hz. (C) Voltage dependence of steady-state activation, normalized so that the activation curve had a maximum value of 1 (assuming fully open channels). (D) Activation time constant. Red data points were estimated from the deactivation protocol (B), while blue data points were estimated using the activation protocol (A). Dashed lines in (C-D) denote the kinetics scheme when corrected for a liquid junction potential of −15 mV for the experimental conditions used for recording Ca^2+^ currents.

The peak current (*I*_*max*_) was recorded for each holding potential in Fig. 8A, and the maximum peak was observed at about 20 mV. By observations, (*I*_*Ca*_) became an outward current when step potentials were increased beyond 70 mV, and based on this we assumed a reversal potential of *E*_*Ca*_ = 70 mV. Similar to what we did for the Na^+^ channel, the maximal conductance (*g*_*max*_) for each holding potential was computed by the equation:

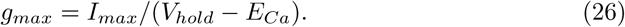

The estimates of *g*_*max*_ for activation are indicated by the crosses in Fig. 8C. The dependency of *g*_*max*_ on *V*_*hold*_ was then fitted by a Boltzmann curve (Eq 23), with *a* = 2 activation variables, and a good fit was obtained with values for *a*, *k* and *V*_*_ as in Table 3.

**Table 3.** Parameters for Ca^2+^ activation. The parameters *p*1-*p*6 are used together with Eq. 25 to yield the time constants for steady state activation (in units of ms). The remaining parameters are used together with Eq. 23 to obtain the steady state activation function. The curves obtained in this way describe the voltage dependence under experimental conditions, and was afterwards corrected with a liquid junction potential of −15 mV (see Fig 2B).

| parameter | activation value | unit |
| --- | --- | --- |
| $p_1$ | -0.128 | $(ms \cdot mV)^{-1}$ |
| $p_2$ | -46.7 | mV |
| $p_3$ | 19.0 | mV |
| $p_4$ | -101.54 | $(ms \cdot mV)^{-1}$ |
| $p_5$ | 535.1 | mV |
| $p_6$ | -60.0 | mV |
| $V_*$ | -6.79 | mV |
| $k$ | 6.57 | mV |
| $a$ | 2 | |
| $V_{lj}$ | -15 | mV |

#### Time constants for *I*_*Ca*_ activation

Assuming two opening gates (*m*), the time constant for *I*_*Ca*_-activation was derived by fitting the function [42]:

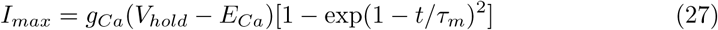

to the response curves in Fig. 8A and B. The activation protocol was used to determine *τ*_*m*_ at high step potentials (from −5 mV and upwards), where the response was not too small and noisy to reveal any clear trend (blue data points in Fig. 8D). The deactivation protocol was used to determine *τ*_*m*_ for lower step potentials (red data points in Fig. 8D). Like for the Na^+^ channel, the voltage dependence of the time constants were fitted using the functional form in eq. 25. Good fits to the data points were obtained with the parameter values in Table 3.

### Adjustments of *I*_*K*_, *I*_*BK*_ and *I*_*SK*_ made in MEDAKA 2

In MEDAKA 2, *I*_*K*_, *I*_*BK*_ and *I*_*SK*_ were adjusted in order to obtain a model whose firing pattern was in closer resemblance with the data, both in terms of the control conditions and under application of paxilline. The adjustments are briefly described below, using the simulations in Fig 9 as a reference.

**Fig 9.**
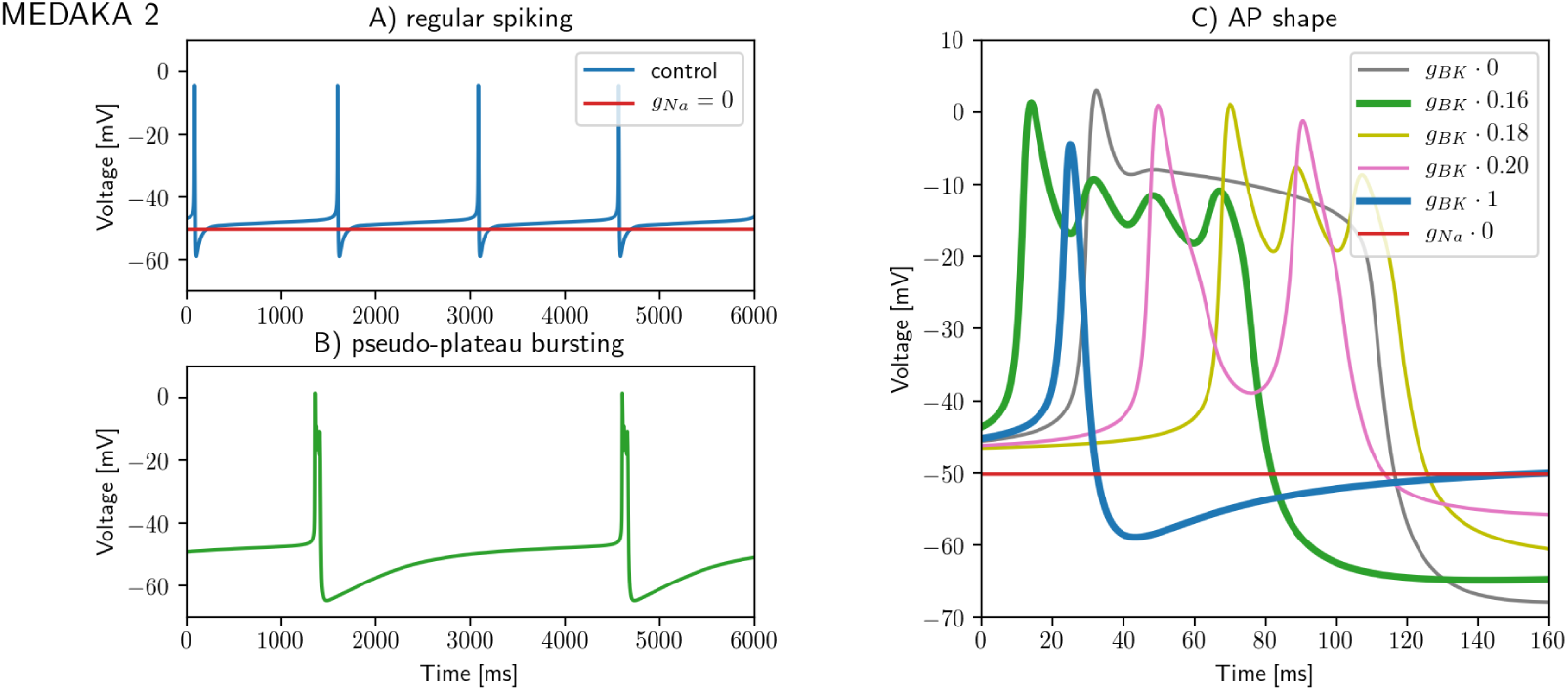
Firing properties of MEDAKA 2. The response properties of MEDAKA 2 were in good agreement with the experimental data in Fig 1. Under the control conditions, the AP firing rate was 0.7 Hz (A). Partial BK-blockage by paxilline made the cell bursty (B), and the burst shape depended on how large fraction of BK channels that were blocked (gray, yellow, pink, and green curves in C). Regular APs peaked at −4.4 mV (blue line in C) and had a duration (taken at half max amplitude between −50 mV and AP peak) of 6.4 ms. These values were within the range of peak and duration values observed in the data. As in the TTX-data (Fig 1C), blockage of *g*_*Na*_ abolished AP generation completely (red line in A).

Firstly, *I*_*BK*_ activation was shifted by 5 mV relative to RAT and MEDAKA 1. This caused *I*_*BK*_ activation to occur closer to the peak, it reduced the effect that blockage of *I*_*BK*_ had on the AP amplitude, and was necessary in order to obtain oscillations during the plateaus following APs (green, yellow and pink curves in Fig 9C).

Secondly, the kinetics of *I*_*K*_ in RAT was based on data from rat lactotropes, where *I*_*K*_ activation had a fast (3.7 ms) and a slow (30 ms) component [54]. The time constant *τ*_*K*_ in RAT was 30 ms, and thus reflected the slow component. For *I*_*K*_ to have an effect on the repolarization of faster *I*_*Na*_-mediated APs, *τ*_*K*_ was reduced to 5 ms in MEDAKA 2, a value closer to the fast component seen in the data [54].

Thirdly, with the maximum value of *g*_*BK*_ in RAT or MEDAKA 1, APs were exceeded by a plateau. *I*_*Ca*_-mediated plateau potentials have not been observed in goldfish [4] or tilapia [5], and was not seen during spontaenous firing in medaka (Fig. 1A-B). To remove the plateaus, *g*_*K*_ and *g*_*BK*_ were increased by factors 1.4 and 4, respectively, compared to RAT and MEDAKA 1, which gave MEDAKA 2 narrow regular APs under control conditions (blue curve in Fig 9C).

Fourthly, the faster repolarization obtained by increasing *g*_*BK*_ and *g*_*K*_ increased the firing rate of MEDAKA 2 to unrealistically high values. Similar effects of *I*_*BK*_ on increasing firing rates have been seen in other systems [39]. A lower and more realistic firing rate was obtained by increasing *g_SK_* by a factor 3.

With the adjustments described above, MEDAKA 2 responded to partial *I*_*BK*_ blockage in a way that resembled the experiments with paxilline application (compare Fig 9C and Fig 1D).

#### Software

Experimental current-clamp data (Fig 1), experimental voltage-clamp data (Figs 1, 6, 7, and 8)) and fitted ion-channel kinetics (Fig 2) were plotted using Matlab (http://se.mathworks.com/). All other plots are made in Python (http://www.python.org).

RAT, MEDAKA 1 and MEDAKA 2 were all implemented using the Python interface for the NEURON simulator [55]. MEDAKA 1 and MEDAKA 2 are original for this work, while RAT was based on a previous model [9].

All simulations (used in Figures 3, 4, 5 and 9) were run for 60,000 ms (although briefer intervals were shown in the figures). The first 10,000 ms were discarded to eliminate initial transients, while the remaining 50,000 ms were used in the uncertainty analysis (Fig 5) and to calculate the burstiness factor (Fig 3). Simulations with noise (Fig 3) were run using a fixed time step *dt* = 0.25 ms, while simulations without noise were run using adaptive time stepping provided by the NEURON simulator.

The sensitivity analysis in Fig 5 was performed by aid of the Python-based toolbox Uncertainpy [34]. The features considered (*Bursting*, *Regular spiking*, and *AP firing* as defined below) were custom made for the analysis in the current work. Uncertainpy was run using polynomial chaos with the point collocation method (the default of Uncertainpy) and a polynomial order of five. The sensitivity analysis was based on calculating Sobol indices. Only the total order Sobol indices were presented in this work. A total order Sobol index quantifies the sensitivity of a feature to a given parameter, accounting for all higher order co-interactions between the parameter and all other parameters (see [56] or the brief overview in Appendix B of [57]).

The models (RAT, MEDAKA 1 and MEDAKA 2), and the code for generating Figures 3, 4, 5 are available for download (doi:10.5281/zenodo.1491552).

### Definitions

Below, the various metrics used throughout this article are defined.

- **AP width:** In control conditions, AP width was defined as the time between the upstroke and downstroke crossings of the voltage midways between −50 mV and the peak potential.
- **Event duration**: A metric proposed by [9], tailored to capture duration of longer events such as bursts. The voltage trace was first normalized so that the minimum voltage was set to 0 and the maximum voltage to 1. The start of an event was defined when the voltage crossed an onset threshold 0.55, and the end of the event was defined when the voltage then crossed a termination threshold 0.45. The duration of the event was the time from onset to termination.
- **Burst**: An AP event with a duration of 60 ms or more was classified as a burst.
- **Burstiness Factor**: The fraction of AP events within a given simulation that were bursts. This metric was most relevant in simulations with noise input (Fig 3). In simulations without noise added, all APs within a given simulation tended to be close to identical, so that the burstiness factor was either 0 or 1 (although there were exceptions).
- ***Isbursting***: In the sensitivity analysis (Fig 5), the feature *Isbursting* was a binary variable that was 1 for simulations that contained bursts, and 0 for simulations that did not.
- ***Isregular***: In the sensitivity analysis, the feature *Isregular* was a binary variable that was 1 for simulations that contained regular APs, and 0 for simulations that did not.
- ***Isfiring***: In the sensitivity analysis, the feature *Isfiring* was a binary variable that was 1 for simulations that contained any APs (regular or bursts) and 0 for simulations that did not.

